# Dissecting transcriptomic signatures of neuronal differentiation and maturation using iPSCs

**DOI:** 10.1101/380758

**Authors:** EE Burke, JG Chenoweth, JH Shin, L Collado-Torres, SK Kim, N Micali, Y Wang, RE Straub, DJ Hoeppner, HY Chen, A Lescure, K Shibbani, GR Hamersky, BN Phan, WS Ulrich, C Valencia, A Jaishankar, AJ Price, A Rajpurohit, SA Semick, R Bürli, JC Barrow, DJ Hiler, SC Page, K Martinowich, TM Hyde, JE Kleinman, KF Berman, JA Apud, AJ Cross, NJ Brandon, DR Weinberger, BJ Maher, RDG McKay, AE Jaffe

## Abstract

Human induced pluripotent stem cells (hiPSCs) are a powerful model of neural differentiation and maturation. We present a hiPSC transcriptomics resource on corticogenesis from 5 iPSC donor and 13 subclonal lines across nine time points over 5 broad conditions: self-renewal, early neuronal differentiation, neural precursor cells (NPCs), assembled rosettes, and differentiated neuronal cells that were validated using electrophysiology. We identified widespread changes in the expression of individual transcript features and their splice variants, gene networks, and global patterns of transcription. We next demonstrated that co-culturing human NPCs with rodent astrocytes resulted in mutually synergistic maturation, and that cell type-specific expression data can be extracted using only sequencing read alignments without potentially disruptive cell sorting. We lastly developed and validated a computational tool to estimate the relative neuronal maturity of iPSC-derived neuronal cultures and human brain tissue, which were maturationally heterogeneous but contained subsets of cells most akin to adult human neurons.

## Introduction

Human induced pluripotent stem cells (hiPSCs) present a unique opportunity to generate and characterize different cell types potentially representative of those in the human brain that may be difficult to ascertain during development or isolate from postmortem tissue. Better characterizing corticogenesis, and identifying subsequent deficits in psychiatric and neurological disorders, has been the focus of much hiPSC and human embryonic stem cell (hESC) research over the past decade (*1*). These cellular models have been especially appealing in neurodevelopmental disorders where direct analyses of neurodevelopmental changes in patients that would be diagnosed with disorders later in life are very challenging (*2*).

Transcriptomics has become an increasingly powerful tool in both brain and stem cell research. Alternative RNA splicing has been revealed to regulate cortical development (*3*). Specific mRNA isoforms also associate with genetic risk for psychiatric disease and their regulation has been modeled in human iPSCs as they differentiate towards neural fates (*4*). Ongoing large-scale efforts use gene expression data from postmortem human brains to further identify molecular mechanisms that mediate genetic risk for psychiatric disease (*5*). However, to date, many large-scale transcriptomics efforts in stem cells with respect to corticogenesis have either focused on mouse ESCs (*6*), or single hESC lines in bulk differentiating cells like CORTECON (WA-09 line) (*7*) or single cells like Close et al. (H1 line) (*8*). Large scale hiPSC resources have focused on the identity and quality of the initial iPSCs without corresponding RNA-seq data following these iPSCs through differentiation, like Kilpinen et al. (301 donors) (*9*), iPSCORE (222 donors) (*10*), and Salomonis et al. (58 iPSC lines) (*11*), which were all subject to extensive quality control steps (*12*).

Here we present the results of a large-scale hiPSC transcriptomics study on corticogenesis from multiple donors and replicate lines across nine time points (days 2, 4, 6, 9, 15, 21, 49, 63, and 77 *in vitro*) that represent defined transitions in differentiation: self renewal, early differentiating cells (accelerated dorsal), neural precursor cells (NPCs), assembled neuroepithelial rosettes (*13*), and more differentiated neuronal cell types (Table 1). We first identified widespread transcriptional changes occurring across the model of corticogenesis including novel alternative transcription, splicing, and intron retention that were largely not influenced by the different genetic backgrounds of the iPSCs. We reprocessed and reanalyzed existing hESC- and iPSC-based resources across both bulk and single cell data in the context of our differentiation signatures which showed similar trajectories of neurogenesis. We then demonstrated *in silico*, that more molecularly mature neuronal cultures are achieved through the addition of rodent astrocytes. We lastly demonstrated that almost half (48%) of the RNA in our neuronal cultures after 8 weeks of differentiation reflected signatures of adult cortical neurons. These data, and software tools for barcoding the maturity of iPSC-derived neuronal cells, are available in a user-friendly web browser (stemcell.libd.org/scb) that can visualize genes and their transcript features across neuronal differentiation and corticogenesis. We anticipate that these data and our approaches will facilitate the identification and experimental interrogation of transcriptional and post-transcriptional regulators of neurodevelopmental disorders.

**Table 1:** Sample and cellular condition information.

|  | Self-renewal | Differentiation time-course |  |  |  | Neuron | Rat astrocyte |
| --- | --- | --- | --- | --- | --- | --- | --- |
|  |  | Accelerated Dorsal | NPC | Rosette | Neuron + Rat astrocyte |  |  |
| **Time point (day)** | 2, 4, 6 | 2, 4, 6, 9 | 15 | 21 | 49, 63, 77 | 77 | 49, 63, 77 |
| **donor_3** | 6 | 8 |  |  |  |  |  |
| **donor_21** | 3 | 12 | 4 | 3 | 10 | 1 |  |
| **donor_66** | 3 | 10 | 3 | 3 | 7 | 1 |  |
| **donor_90** | 3 | 10 | 2 | 1 | 7 | 1 |  |
| **donor_165** | 3 | 14 | 3 | 2 | 7 | 1 |  |
| **Rat** |  |  |  |  |  |  | 3 |
| **Total** | 18 | 54 | 12 | 9 | 31 | 4 | 3 |

See: Table1\_sample\_information.xlsx

## Results

### Differentiating hiPSCs to electrophvsioloqically-active neuronal cultures

We reprogrammed fibroblasts from the underarm biopsies of 14 donors/subjects creating a median 5.5 iPSC lines (interquartile range: 3-7) per subject that were free from karyotyping abnormalities (see Methods). Fluidigm-based qPCR expression profiling confirmed the loss of *FAP* (Fibroblast Activation Protein Alpha) expression across all iPSC lines (Figure S1A, p < 2.2×10-16) and the gain of *NANOG* expression (Figure S1B, p=1.87×10-13). We ultimately differentiated iPSCs from five donors and fourteen total lines towards a neural stem cell specification, followed by cortical neural progenitor cell (NPC) differentiation and expansion, followed by neural differentiation/maturation (see *Methods*). RNA-seq of these lines confirmed the expected temporal behavior of canonical marker genes in thirteen of the lines, including the loss of pluripotency gene *POU5F1/OCT4* (Figure 1A) and gain of *HES5* (Figure 1B) through NPC differentiation, and the gain of *SLC17A6/VGLUT2* expression through neural maturation (Figure 1C). High-content imaging confirmed the selforganization of NPCs into neuroepithelial rosettes (*13*) (Figure 1D) and electrophysiological measures like capacitance and membrane resistance taken at 49, 63, and 77 days *in vitro* (DIV, corresponding to 4, 6, and 8 weeks following NPC expansion) show electrophysiological maturation (*14*) (Figures 1E, 1F). This highlights the ability of our protocol to create neuronal cell lines that display hallmark signatures of neuronal differentiation and are electrophysiologically active.

**Figure 1:**
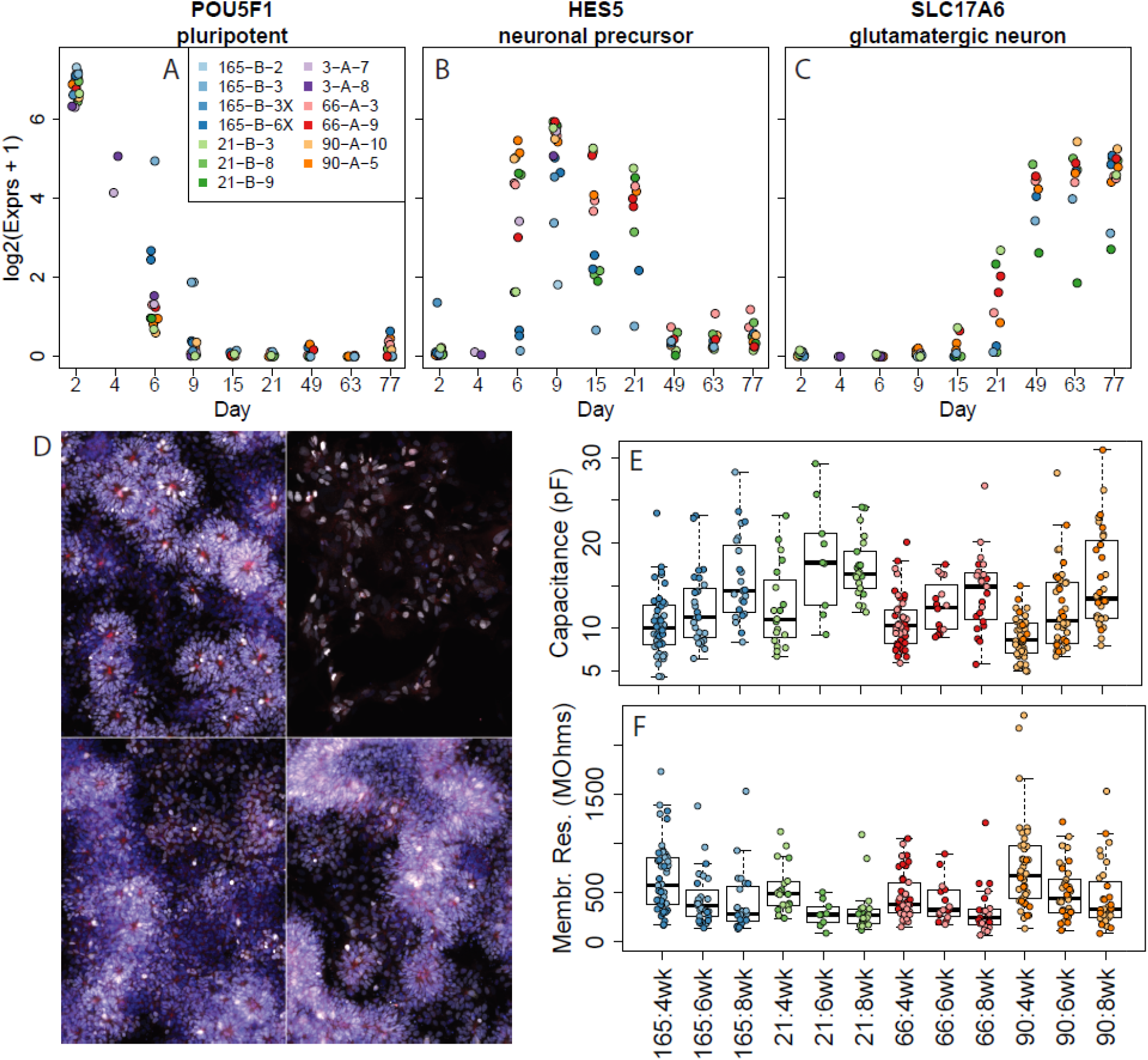
Differentiating hiPSCs follow expected trajectories of neuronal development. Normalized expression levels from RNA-seq showing the expected temporal behavior of canonical marker genes through differentiation: (A) the loss of pluripotency gene *POU5F1/OCT4*, (B) the expression of *HES5* through NPC differentiation, and (C) the gain of *SLC17A6/VGLUT2* through neural maturation. (D) Presence of self-aggregating neural rosettes using representative images from one subclonal line across four donors. Lines clockwise from top left: 66-A-9, 21-B-9, 165-B-3, and 90-A-10. Blue - DAPI; Red - ZO-1; White - OTX2. Electrophysiology measurements across neuronal maturation show(E) increasing capacitance and (F) decreasing membrane resistance.

### Global transcriptional signatures of differentiating and maturing neural cells

We first sought to transcriptionally characterize this iPSC model of corticogenesis across five conditions: self-renewal, dorsal fate specification, NPCs, self-organized rosettes, and maturing neural cells. We therefore performed stranded total RNA-seq following a ribosomal depletion on a total of 165 samples, sampling from nine time points across five donors and the five conditions above and a series of technical samples (see *Methods*). While samples were sequenced across 11 flow cells, we repeated three RNAs on seven flow cells, and the gene expression profiles of these samples clustered by sample (Figure S2A), whereas synthetic ERCC spike-ins (performed in each library) clustered more by flow cell (Figure S2B). Following RNA-seq, we obtained high proportions of exonic reads to Gencode v25 even among the neuronal cells co-cultured with rat astrocytes (Figure S2C). We dropped the 6 samples from one line (165-B-8X) that differentiated more slowly than other lines (using *NANOG* expression as a proxy, Figure S2D). Lastly, we dropped 5 samples with sample identity mismatches using called coding variants matched to existing microarray-based genotype calls (Figure S3).

We first confirmed the representativeness of our iPSC cell lines and subsequent differentiation with the recently published ScoreCard reference data (*15*) (Figure S4A) — our self-renewal/iPSC lines showed mean 98.1% pluripotency identity (standard deviation (SD)=1.5%) which significantly decreased through differentiation (Figure S4B, p<2.2×10-16). We then characterized the global landscape of transcriptional changes accompanying these differentiating and maturing cells using principal component analysis (PCA) of gene expression levels. The largest component of variability in the data represented corticogenesis (PC1, 52.4% of variance explained), while the second PC (14.3% of variance explained) further separated those samples in the NPC stage from self-renewing and neuronal cells (Figure 2A). Both of these components of variability were highly conserved in reprocessed data from the CORTECON hESC time-course dataset (Figure 2A, Figure S5, PC1 p=3.49e-9, PC2 p=1.65e-8) even though these cell lines were ESCs differentiated, processed, and sequenced in different labs. We further identified global similarity between our differentiating and maturing cells to recently available single and pooled cell-level data from Close et al. (*8*), reprocessed using the same pipeline (Figure S6, see *Methods*).

**Figure 2:**
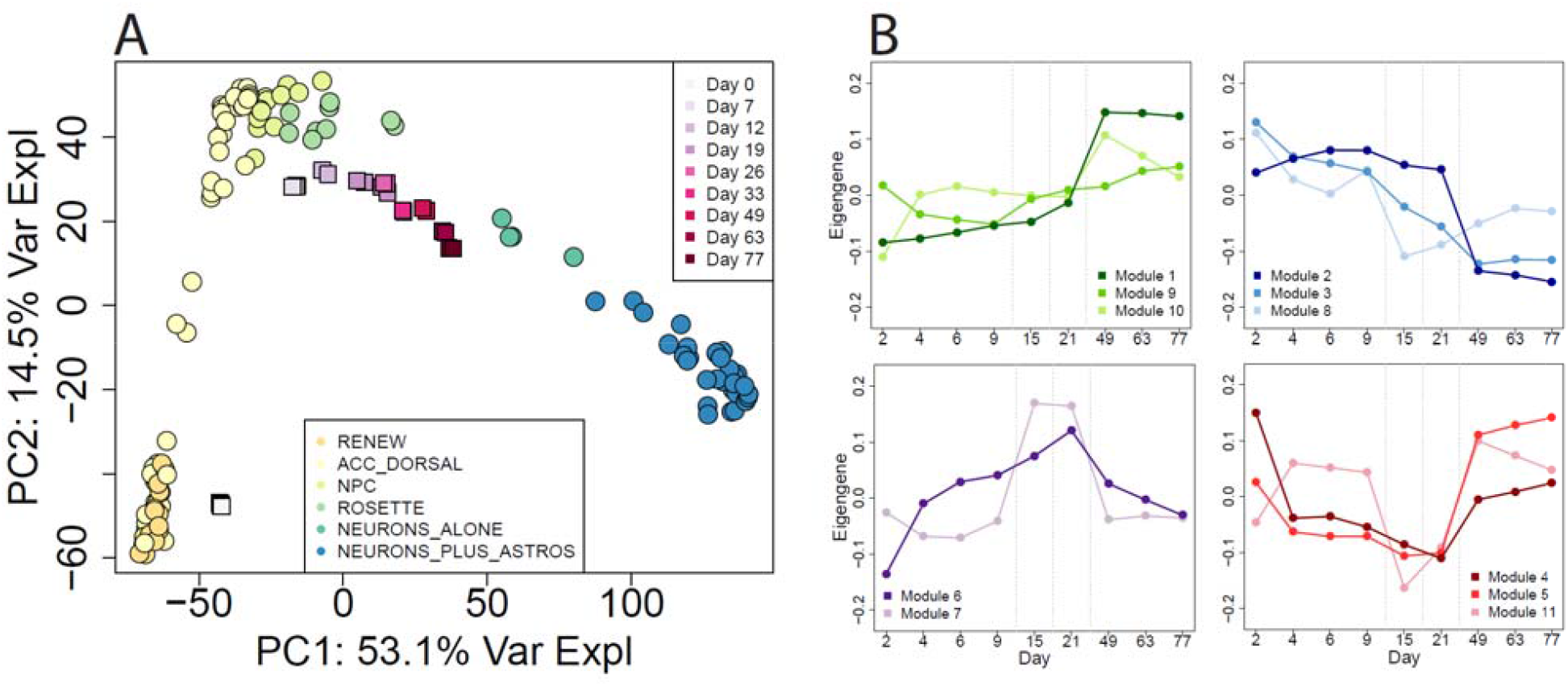
Global expression comparison to CORTECON. (A) PCA of gene expression levels showing PC1 representing corticogenesis and PC2 separating samples in the NPC stage from self-renewing and neuronal cells, as well as the conservation of these components of variability in the CORTECON dataset. (B) Eigengenes of the eleven WGCNA modules created from the RNA-seq data, grouped by dynamic expression pattern: genes that turn on in mature neurons, those that turn off in mature neurons (related to loss of pluripotency), those that rise in NPCs then fall, and those that fall in NPCs then rise again in neurons.

We then performed weighted gene co-expression network analysis (WGCNA) (*16*) to identify more dynamic and robust patterns of expression across neural differentiation and maturation. We identified eleven signed co-expression modules across the 25,466 expressed genes (Figure 2B, with 3,284 genes in grey/unassigned module). These modules reflected known stages of differentiation, which were confirmed using gene set enrichment analysis (Table S1), including genes related to loss of pluripotency (Module 3), rising in NPCs (Module 7) and also turning on (Modules 1 and 5) and off (Module 2) in neuronal cultures. We calculated the corresponding eigengenes in these modules in the reprocessed CORTECON dataset, which showed replication for many of the same temporal patterns observed in the discovery dataset (Figure S7). These differentiating cells therefore show analogous global expression patterns as other previously published datasets of corticogenesis (*7, 8*).

### Alternative splicing and previously un-annotated expression of differentiating neural cells

We next tested for developmental regulation of individual genes and their transcript features among the 106 time-course samples (excluding 18 self-renewing lines and 4 neuronal lines that were not cultured on rodent astrocytes, Table 1) using statistical modeling (see *Methods*). By partitioning gene expression variability into different components, we demonstrated that differentiation (condition) explains much more variability in expression than donor/genome or sub-clonal line for the majority of expressed genes (19475 genes, 76.4%, Figure 3A). The majority of expressed genes changed in expression (false discovery rate (FDR)<0.01) across differentiation and maturation (20,220 genes, 79.4%), including 9067 genes differentially expressed between accelerated dorsal differentiation and NPCs, 1994 genes between NPCs and rosettes, and 12,951 genes between rosettes and neuronal cultures. Across the time-course we also found widespread differential expression at the exon (70.5% of unique genes), exon-exon splice junction (66.2%), and full-length transcript summarizations (72.1%) (Figure 3B, 3C, Table S2). We have created a user-friendly database to visualize these feature-level expression across neuronal differentiation, available at stemcell.libd.org/scb.

**Figure 3:**
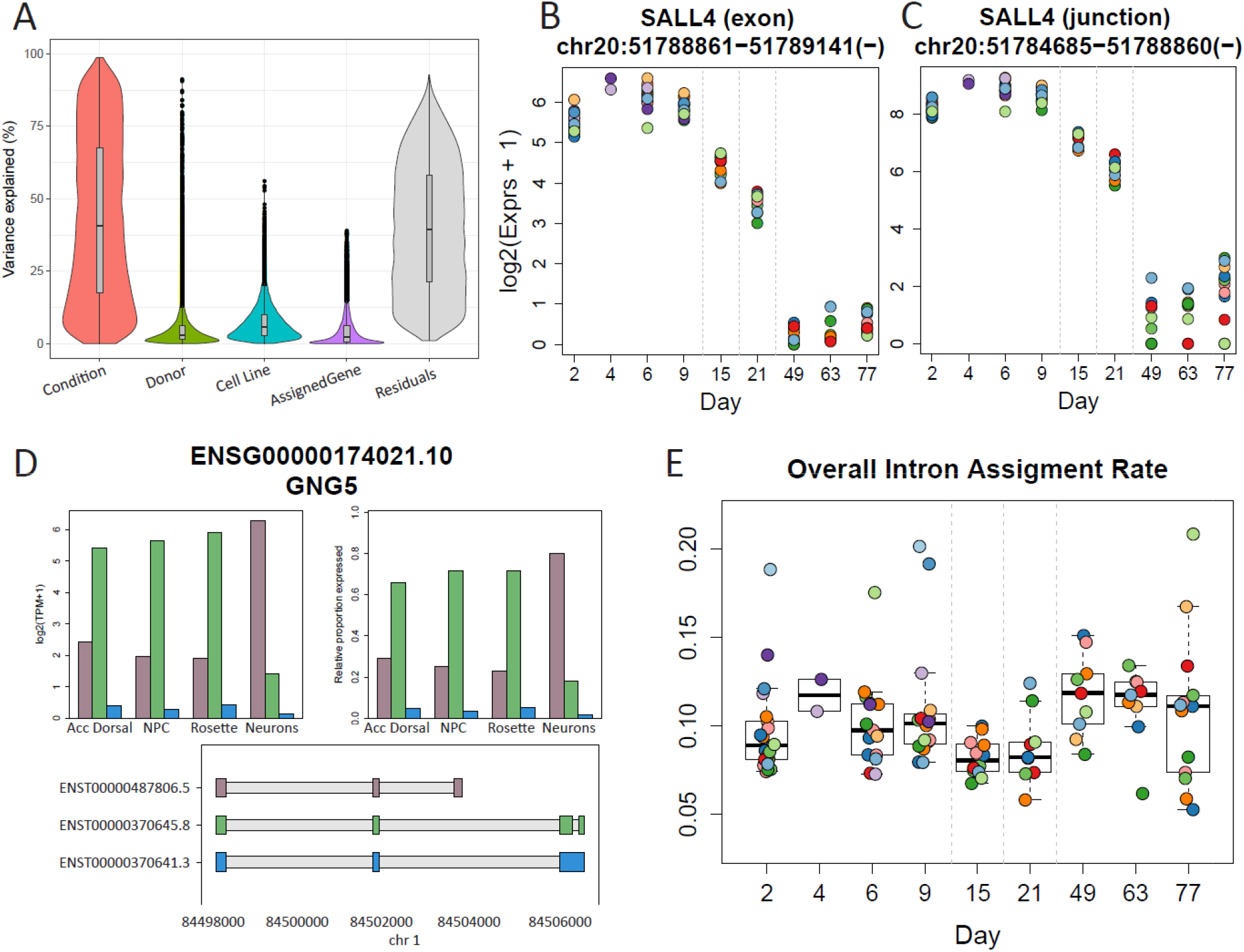
Feature-level differential expression. (A) The contribution of variance in expression models on the gene level. (B) An example of a differentially expressed exon across conditions from the *SALL4* gene, which is thought to play a role in the development of motor neurons, as well as expression of the neighboring exon-exon junction (C). (D) An example of a gene with shifting major isoform usage, with one transcript highly expressed in the first three conditions, shifting to a second highly expressed transcript in the neuron condition. (E) Percent of aligned reads assigned to intronic sequences across all samples and timepoints, with a significant overall gain in intronic assignment rate between NPCs/rosettes (days 15 and 21) and differentiated neurons (days 49-77) in line with previous research.

Interestingly, we found a large number of previously un-annotated junctions that were differentially expressed across differentiation, including 7298 exon-skipping events and 15,002 alternative exonic boundaries (Table S3), a much larger number than previously reported in smaller studies of hESC differentiation (*3*). As many of these isoforms could represent cell type- or developmentally-specific events, we sought to more fully characterize how differentiating genomes use multiple transcript isoforms of the same genes differently. We identified the highest expressed transcript across samples of each cell condition, and found that 7,856 out of 25,466 expressed genes (30.8%) showed change in the most abundant transcript isoform across the four conditions. The highest percentage shift occurred between rosettes and neuronal cells, where 6,459 genes had major isoform shifts. Of the 12,951 genes we found to be differentially expressed between rosette and neuron, 3,998 had a shifting isoform (Table S4, Figure 3D). These genes highlight examples of diverse developmentally-regulated neuronal cell differentiation stages using the same gene in different ways across corticogenesis.

In an additional analysis related to splicing, we looked into the increase of intron retention (IR), which has previously been shown to play an important role in differentiation (*17*). We identified an overall gain in the percentage of aligned reads assigned to introns between our NPCs/rosettes and differentiated neuronal cells (Figure 3E, p=0.002), in line with previous research describing intron retention as a mechanism of rapid gene regulation in response to neuronal activity (*18*). Additionally looking at IR ratios, gene ontology (GO) analysis on the list of 1329 genes with significantly (FDR<0.001) increasing IR ratios through differentiation showed strongest enrichment for neuronal biological process including synaptic signaling (GO:0099536, p=9.23e-7, adjusted p=6.9e-4) and neuron development (GO:0048666, p=9.87e-7, adjusted p=6.9e-4) (Figure S8). These analyses confirm the extensive transcriptional changes occurring during neural differentiation and neuronal maturation among individual transcript classes.

### Co-culturing neuronal precursors on astrocytes accelerates maturation

Given the relatively diminished progenitor-like signature of neuronal cells that were co-cultured at the NPC stage with rat astrocytes compared to neuronal cells cultured alone (PC2 in Figure 2A), we sought to more fully characterize the transcriptional effects of astrocyte co-culturing. We first demonstrated that we could accurately separate the expression data from human neuronal cells and rat astrocytes from co-cultured cells using RNA-seq read alignment, as an “*in silico*” RNA sorting technique. We analyzed RNA-seq data from rat astrocytes and human neuronal cells alone and found little cross-species mapping — human neuronal cells alone had low alignment rates to the rat (rn6) genome (mean=16.6%, SD=4.5%) and rat astrocytes alone had low alignment rates to the human (hg38) genome (mean=10.1%, SD=2.3%) (Figure S9). We next re-mapped those rodent reads that aligned to hg38 back to rn6 and those human reads that aligned to rn6 back to hg38 and found two sets of three highly expressed genes that contributed to the majority of reads that mapped across species (Table S5). These data suggest that expression profiles of human neuronal cell classes can be computationally separated from rodent astrocytes, eliminating the need to perform flow cytometry which can introduce expression changes in RNAs (*19*).

We then compared the human gene expression profiles between four neuronal lines cultured alone and seven of the same lines co-cultured with rodent astrocytes at week 8, and identified 3214 genes differentially expressed (at FDR < 0.05) between the two groups (Figure 4A, Table S6). We performed GO analyses on the sets of up- and down-regulated differentially expressed genes, and found very significant enrichment of genes related to transporter activity, ion channels, and their activity among co-cultured neuronal cells plus astrocytes (Figure 4B, Table S7). The up-regulated genes were further enriched for being localized in the ion channel complex (GO:0034702, p=1.36e-25, adjusted p=6.86e-23), post-synapse (GO:0098794, p=1.35e-24, adjusted p=3.4e-22), and axon (GO:0030424, p=1.68e-21, adjusted p=1.65e-19). These results corroborate previous observations that co-culturing with astrocytes produces cultures with more mature neuronal cell types (*20*). We next examined the IR ratios of the lines cultured alone and found their IR ratios to be similar to the less mature NPCs/rosettes (p=0.68) and lower than those of the neuronal samples co-cultured with astrocytes (p=0.036). Lastly, we explored these gene expression associations using electrophysiology, and showed increased neuronal maturation from weeks 4 to 8 of co-culturing with astrocytes, through both increasing capacitance (p < 2.2e-16, Figure 1E) and decreasing membrane resistance (p=2.15e-14, Figure 1F).

**Figure 4:**
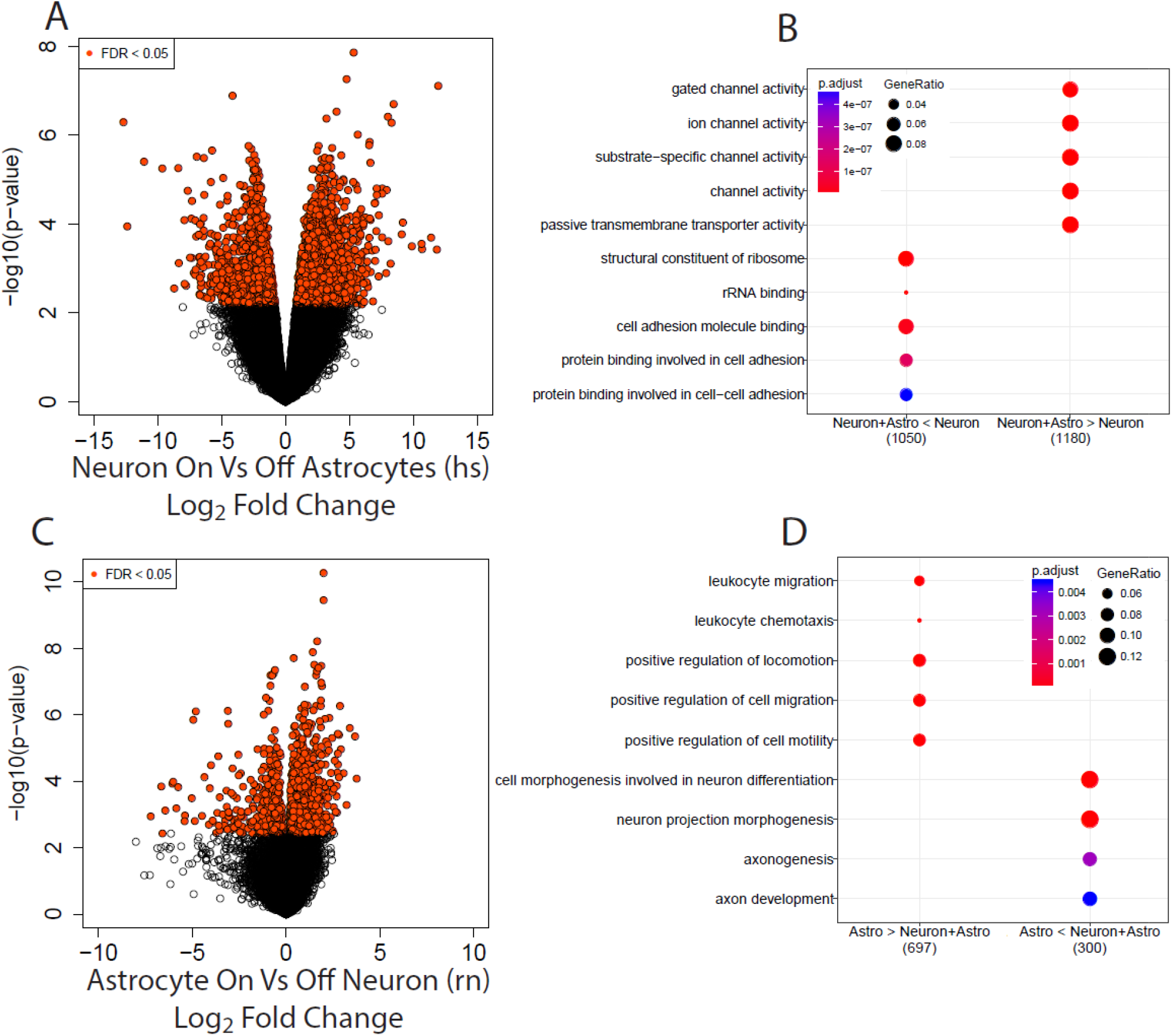
Astrocyte co-culturing. (A) Volcano plot showing the differential expression between human neurons cultured alone and human neurons co-cultured with rat astrocytes, as well as the enrichment (B) of the 3214 genes differentially expressed at FDR < 5%, split by direction of change. (C-D) Analogous results of differential expression analysis quantified against the rat genome, comparing rat astrocytes alone with rat astrocytes co-cultured with human neurons.

As a secondary analysis, we turned to the rodent astrocytes (quantified against the rat transcriptome) and asked whether co-culturing these with human neuronal cells altered their transcriptomes. We found 1329 rodent genes differentially expressed between astrocytes cocultured with neuronal cells compared to astrocytes alone (at FDR < 0.05, Figure 4C, 4D), and those genes more highly expressed when co-cultured were strongly enriched for neuron projection morphogenesis (GO:0048812, p=9.78e-9, adjusted p=1.66e-05) and axon development (GO:0061564, p=5.19e-6, adjusted p=4.4e-3). These results therefore suggest increased synergistic maturation of both neuronal cells and astrocytes when co-cultured together.

### RNA deconvolution quantifies subpopulations of mature cortical neuronal cells

Given the expression trajectories and physiological activity among the 8 week neuronal cells, we sought to more fully characterize the underlying cellular composition of these cultures. Previous computational approaches have focused on global analyses determining the most representative time point in brain development for iPSCs and organoids (*21*). Here we instead developed a strategy to quantify the fraction of RNAs from ten different developmental, prenatal, and postnatal neural cell types. We re-processed and jointly interrogated microfluidics-based single cell RNA-seq datasets from iPSCs and NPCs, and fetal quiescent and replicating neurons, and adult neurons, astrocytes, oligodendrocytes and their progenitors, microglia, and endothelial cells (*22, 23*). These single cell data were selected to be more directly comparable to RNA-seq of bulk cells, including the quantification of the entire gene body (rather than 3’ read counting) and fresh brain tissue (rather than frozen tissue, that results in ruptured cell membranes and the need for subsequent sequencing of nuclear, rather than total, RNA). We identified a set of 228 genes that could transcriptionally distinguish each of these ten cell classes from the others using feature-selection strategies previously described for cellular deconvolution with DNA methylation data (*24*), standardized the expression to reduce technical effects across studies (i.e. created Z-scores), performed regression on these RNA profiles to estimate the mean RNA levels for each cell class, and then implemented the quadratic programming-based approach of (*25*) to perform RNA deconvolution (see Methods).

We describe the algorithm in Figure 5 using data from a subset of genes (to aid visualization). Figure 5A displays the mean standardized expression levels for each cell class for the 131 genes that distinguish iPSCs, NPCs, fetal replicating neurons, fetal quiescent neurons, adult neurons, and adult endothelial cells (vertical lines). Our standardized data across these 131 genes is shown in Figure 5B, which largely depicts blocks of decreased expression of iPSC and NPC genes, and increased expressed of fetal and adult neuronal genes. The expression levels of two adult neuronal genes selected by the algorithm (*SNAP25* and *SCN2A*) across the single cell reference and our bulk data are shown in Figures 5C — identical boxplots can be made for all genes in this signature (Figure S10). The RNA deconvolution algorithm estimates how similar the expression profile of each sample is to each of the ten reference profiles across these genes, and computes the RNA fraction of each cell class — the shifts of these fractions across differentiation and maturation are shown in Figure 5D.

**Figure 5:**
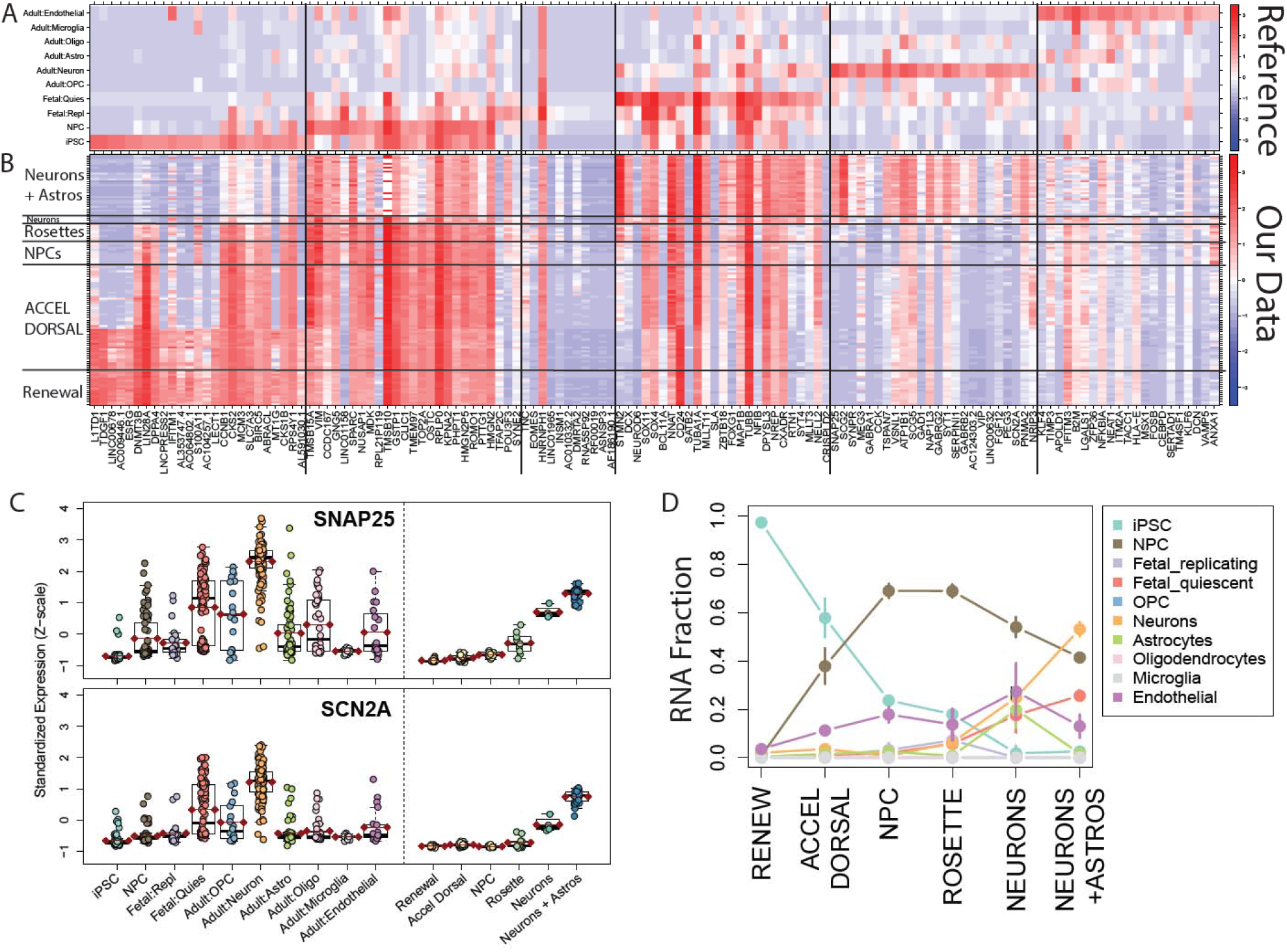
Deconvoluting RNA from underlying cell types across differentiation. (A) The mean standardized expression in the single cell reference data of the 131 genes that were found to distinguish iPSCs (25 genes), NPCs (25 genes), fetal replicating neurons (11 genes), fetal quiescent neurons (25 genes), adult neurons (24 genes), and adult endothelial cells (21 genes). (B) The mean standardized expression of our time-course data across these 131 genes, showing expected higher expression of iPSC genes in the earlier time-course samples, and higher adult neuronal genes in the samples of neurons on astrocytes. (C) Boxplots of the standardized expression in both the reference and time-course data at the single cell level of two genes, *SNAP25* and *SCN2A*, that distinguish adult neurons. (D) The RNA fraction of cell types of our bulk data estimated by the deconvolution algorithm, showing the fall of iPSCs and the rise of fetal quiescent and adult neurons.

More formally, we observed the loss of RNA expression signatures from iPSCs (p=1.3e-14), the rise and fall of RNA expression signatures from NPCs (with a similar pattern as the PC2 of the gene expression data in Figure 2A), and rise of RNA expression signatures from fetal quiescent (p=3.4e-33) and adult (p=6.4e-32) neurons (Figure 5D). We also observed significantly increased RNA fractions of adult-like neuronal cells when plated on (48.9%) versus off rodent astrocytes (23.4%, Figure 5D). The mixture of estimated RNA fractions from neuronal classes from diverse developmental stages highlights the heterogeneity of maturation states from iPSC-derived neuronal cells. However, this approach further demonstrates that a subpopulation of cells in these neuronal cultures are more transcriptionally akin to adult neurons than generally thought, and also provides a computational tool for transcriptionally assessing the relative maturity of differentiated neurons from iPSCs (via the RNA fraction from adult neurons) across independent datasets and experiments. We do emphasize that this algorithm, when applied to expression data, only estimates the RNA fraction of each cell class, and explicitly not the proportion of cells present in the culture — the exact link between RNA proportion and cellular proportion is unclear. For example, if more mature neurons are larger and more transcriptionally active, they would contain more RNA than other less mature neuronal types.

We performed several additional analyses to independently assess this RNA deconvolution tool, particularly the larger fraction of RNA derived from more mature neuronal cells than previously reported in the literature (*21*). First, we applied this RNA deconvolution approach to a large RNA-seq dataset from postmortem human brain tissue (*26*), and found largely expected developmentally regulated signals. We observed loss of fetal quiescent neurons (p<1e-100) and NPCs (p=6.5e-91) and the rise of adult neurons (p=5.0e-39), and oligodendrocytes (p=7.4e-54), primarily at the transition between pre- and post-natal life (Figure S11). The estimated RNA fraction of neuronal RNA in the adult samples (mean=61.9%) was almost twice as high as previous cell count-based approaches, including cytometry-based fractions of NeuN+ cells (mean ~33%) (*27*) and DNA methylation-based deconvolution of frontal cortex (27.9%) (*28*), but was lower than other RNA-based deconvolution strategies applied to DLPFC RNA-seq data (mean=80%) (*29*). Second, we designed and implemented an orthogonal RNA deconvolution model using bulk RNA-seq data from iPSCs and the BrainSpan project to estimate the RNA fractions from eight developmental stages (iPSC, early-, mid- and late-fetal cortex, and infant, child, teen, and adult cortex) using 169 genes (see Methods). We again found loss of pluripotency, rise and fall of the early-fetal signature, and then rise of both mid-fetal but also adult cortical signatures (Figure S12). This developmental stage-based deconvolution further demonstrated the emergence of a more adult-like RNA signature in neuronal cells co-cultured on astrocytes.

### Characterizing the cellular landscapes of other experimental systems and datasets

We assessed the robustness of this RNA deconvolution strategy for assessing neuronal maturity across a series of publicly available datasets on human brain and neuronal differentiation *in vitro*. Using data from the BrainSpan project (*30*), we confirmed the shift from fetal quiescent to mature adult neurons in homogenate RNA-seq data (Figure 6A) and the shift from replicating to quiescent fetal neurons in 5-20 post-conception weeks (Figure 6B) and independent 33-91 post-conception day (corresponding to 5.5-13 PCW, Figure 6C) fetal neocortex single-cell RNA-seq datasets.

**Figure 6:**
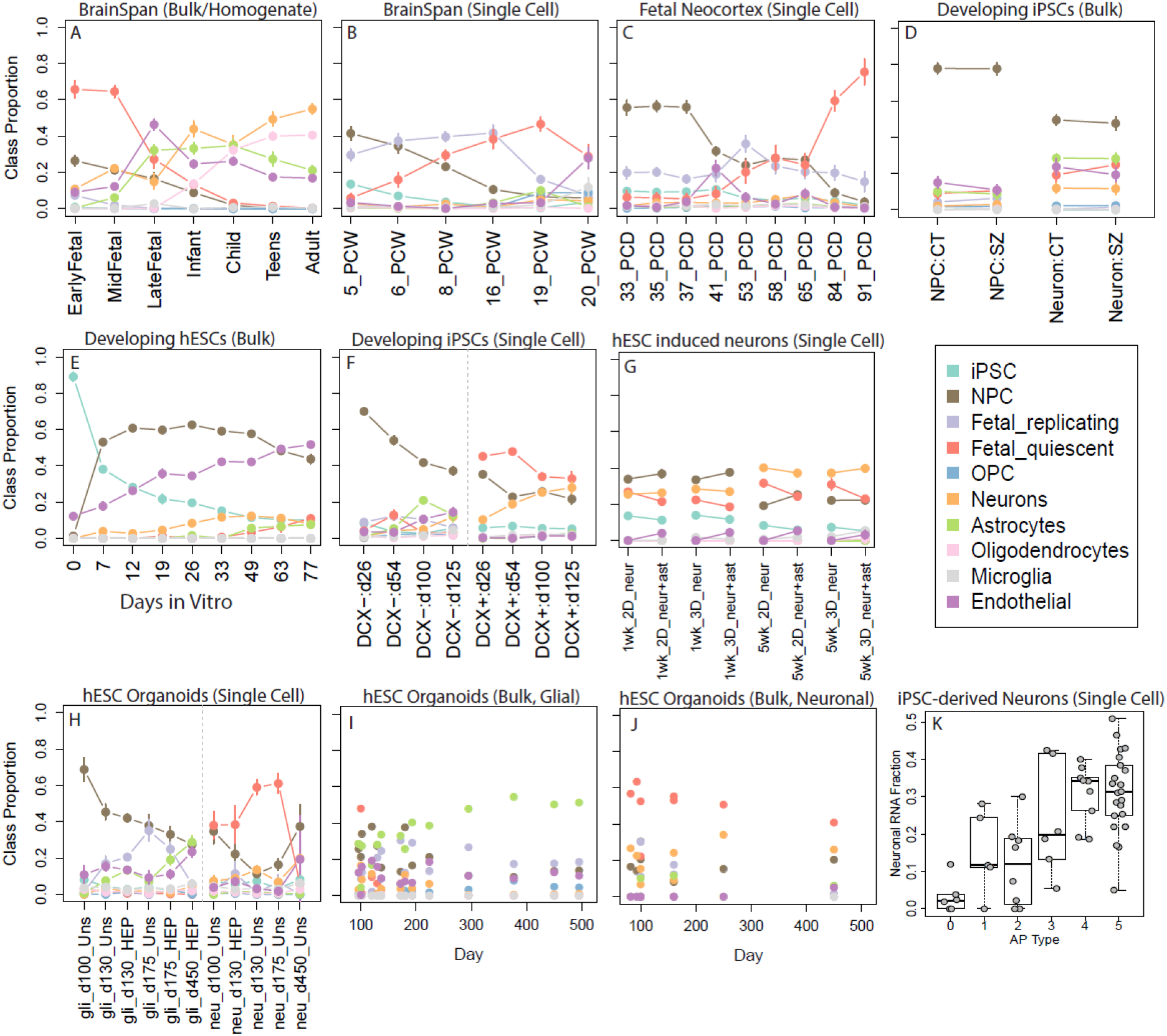
Deconvoluting RNA fractions from publicly available RNA-seq data. We assessed the robustness of our deconvolution algorithm by applying it to multiple RNA-seq datasets. (A-C) Bulk and single cell samples from the BrainSpan project showed expected cellular trajectories for both pre- and post-natal samples. (D) Schizophrenia and neurotypical samples showed similar loss of NPCs and rise of neurons and astrocytes through differentiation. (E) In CORTECON data we found a loss of iPSCs and a rise in the endothelial signature — which we hypothesize represents an immature astrocytic signature. (F) The differing RNA fractions of DCX+ and DCX-single cell populations include a sharper rise of adult neurons in the DCX+ samples. (G) The cell signatures of 2D and 3D neurons were similar, showing a higher NPC fraction at 1 week, and higher adult neuronal fraction at 5 weeks of differentiation. (H-J) The cell signatures of iPSC-derived organoids in single cell and bulk samples using different selection strategies. (K) Samples with RNA-seq combined with electrophysiology data suggest a positive relationship between neuronal RNA fraction and activity state, as well as that less mature astrocytes (coded as AP Type 0) are classified with an endothelial cell signature (Figure S12B).

Within bulk RNA-seq data from differentiating neuronal cells (*31*), we found increases in the proportion of neurons and astrocytes through differentiation in human iPSCs in a recent schizophrenia and control collection (Figure 6D). This manuscript had reported a residual fibroblastlike signature which was not supported by applying this deconvolution to pure fibroblast and iPSC data (Figure S13A). In data from CORTECON (*7*), we further found a larger proportion of endothelial — which we hypothesize could represent an immature astrocyte signature (see below) — and NPCs, and a lack of fetal or adult neuronal classes (Figure 6E). These less mature cell states — possibly due to the CORTECON protocol not including astrocyte co-culturing — were in line with our global PC analysis in Figure 2A. Analyses in single cell differentiation datasets further revealed stark differences in the underlying RNA fractions across development, including between DCX- and DCX+ cell populations, in the shift from RNAs from NPCs to more mature neurons in DCX+ populations (*8*) (Figure 6F).

Other experimental systems and approaches are becoming popular tools in parallel to classical 2D culture conditions, enabling us to make comparisons to both 3D cultures and human organoids. First, we found similar RNA fractions comparing 2D versus 3D directly induced neurons, both on versus off astrocytes, at both 1 and 5 weeks after differentiation (*32*) (Figure 6G). This suggests 3D cultures may not generate more transcriptionally mature cells than standard 2D cultures, at least in an induced neuron protocol. We also profiled the RNA fractions in iPSC-derived organoids over longer periods of differentiation (*33*), and found expected RNA distributions within neuronal and glial cellular subtypes, including the rise of RNA from astrocytes in HEPACAM-selected cells (Figure 6H,I) and RNA from fetal quiescent neurons in neuronal populations (Figure 6J).

We lastly sought to more functionally validate the RNA fraction originating from the adult neuron class using Patch-seq with paired electrophysiology data (*34*). We found a significant positive relationship between the activity states (from 1 to 5, excluding astrocytes, coded as 0) of iPSC-derived neuronal cells and the estimated RNA fraction from neurons (Figure 6K, p=3.44e-5). These data also suggested that the endothelial RNA fraction associated with less mature astrocyte cells (~20% estimate, Figure S13B), is in line with the CORTECON data above. These results from diverse RNA-seq datasets from a variety of experimental approaches further suggest this transcriptional deconvolution strategy can be a robust tool for evaluating the cellular maturity of model systems.

## Discussion

Here we extensively characterized the transcriptomes of human iPS cells as they traverse distinct neurodevelopmental transitions defined by morphological and functional features. We used thirteen independent cell lines derived from five donors to understand the relevance of these cellular systems to model human brain development. We identified widespread transcript-specific changes in the expression of genes across differentiation and neuronal maturation at varying scales, including globally, among co-expressed genes, and among individual transcript features which were assessed for replication in independent neuronal differentiation datasets. It will be important to relate these data to transcriptional isoforms that associate with risk for neurodevelopmental disorders to build faithful *in vitro* models for experimental applications.

We also focused on the ability to assess neuronal maturity in our paradigm. This was approached both experimentally through use of rodent astrocyte co-cultures, and computationally through the use of two related deconvolution implementations to determine the brain stage and cellular identities of differentiating cells that we applied to ten independent datasets across thousands of samples and cells. While previous reports have demonstrated the enhanced neuronal maturity of co-culturing NPCs with astrocytes during differentiation both electrophysiologically and transcriptionally (*35–37*), few studies, to our knowledge, have generated cell type-specific gene expression profiles without the use of cell sorting. Here we show that the functional consequences of co-culturing human NPCs with rodent astrocytes can be detected *in silico* by leveraging the mappability of longer sequencing reads across different species. The resulting read alignments to each species’ genome can largely recapitulate the cell type-specific expression profiles of neurons and astrocytes. These crossspecies analyses were used to show the synergistic effect on both neuron and astrocyte maturation when the two cell types were co-cultured.

We have also leveraged tools previously developed for estimating cell type composition profiles from DNA methylation data to benchmark the developmental and cellular landscapes of human iPS cultures as they differentiate towards neural fates. We show that 8 week neuronal cultures, particularly those co-cultured on rodent astrocytes, are a diverse mixture of cells at varying maturities and cellular states. While the majority of cellular and developmental classes are consistent with the prenatal brain as previously reported using microarray-based profiling (*21*), we have identified subsets of more mature cells also present in our cultures that model later developmental stages. These cells presumably account for those with evidence of more mature phenotypes in our electrophysiology assays. The two references profiles and subsequent regression calibration-based tools for deconvoluting the relative RNA contributions of cell stages and classes can be easily utilized by researchers using RNA-seq data to ensure more comparable case and control lines for discovering molecular phenotypes. This approach will also be increasingly valuable to assess organoids and complex three dimensional stem cell models under development (*38*).

Recent advances in single cell analysis are being leveraged to comprehensively define the human central nervous system. The BRAIN Initiative has effectively used single cell transcriptomics and epigenetics to characterize cellular populations in the developing brain (*39*). We anticipate that the tools developed in our study will complement those efforts that depend on cellular dissociation and selection that largely restrict data to gene-level and nuclear expression. Future development of mathematical and computational approaches to relate datasets and enhance sparse cell-level insight will be valuable towards understanding brain health and disease.

## Methods

### Clinical Fibroblasts

Skin fibroblasts, taken in a superficial circular incision (3mm in diameter) in the mesial aspect of the upper arm, were cultured after informed consent from neurotypical volunteer subjects who were participants in the Sibling Study of Schizophrenia at the National Institute of Mental Health in the Clinical Brain Disorders Branch (NIMH, protocol 95M0150, NCT00001486, Annual Report number: ZIA MH002942053, DRW PI) with additional support from the Clinical Translational Neuroscience Branch, NIMH (KFB PI). All subjects were extensively screened with obtaining medical, psychiatric and neurological histories, physical examinations, MRI scans, and genome wide genotyping to rule out diagnosable clinical disorders.

### iPS cell line derivation and culture

Human iPS cell lines were generated using the Stemgent mRNA reprogramming kit (00-0071) and the Stemgent microRNA Booster kit (00-0073) with modifications (*4*). Briefly, human fibroblasts were seeded with 50,000 cells in individual wells of 6-well plate coated with Matrigel in DMEM media + 10% FBS and 2mM L-glutamine. The next day (day 1), the media was changed with Pluriton human NUFF conditioned media with 300ng/ml B18R protein. On days 1 and 5, the microRNA booster kit was used with the StemFect RNA transfection reagent kit from Stemgent to enhance reprogramming. On days 2-12, the OSKML RNAs were transfected. The mRNA reprogramming process was performed in 37oC, 5% O2 and CO2 incubator. Individual colonies were picked and expanded on irradiated mouse embryonic fibroblasts in DMEM-F12 (Invitrogen), 20% knockout serum replacement (KSR), 5 ng ml^−1^ FGF2 (R&D Systems), 0.1 mM 2-mercaptoethanol (Sigma), 2 mM L-glutamine, and 1× non-essential amino acids (both from Invitrogen). Putative iPS cell lines were subjected to karyotype analysis (Cell Line Genetics) and molecular analysis prior to the generation of feeder-free working cell banks. Confirmation of known pluripotency genes and silencing of fibroblast enriched genes (*40*) in reprogrammed cells was measured using the Fluidigm BioMark System and TaqMan probes according to the manufacturer’s protocol. Only cell lines with a normal chromosomal complement were chosen to generate cell banks for this study. For maintenance of banked hiPSCs in feeder-free conditions, cells were dissociated to single cell populations with accutase (A11105, Life technologies), plated at a density of 1×10^6^ cells in a Matrigel (BD)-coated 6-well plate and cultured with mTeSR1 (Stem Cell Technologies, #05850) (*41*). The cells were cultured with 5 mM Y27632, ROCK inhibitor (Y0503, Sigma-Aldrich) to increase the single cell survival upon dissociation. At 24 hours after plating, Y27632 were removed from the medium and cells were cultured for another 4 days before the next passaging.

### Neural Differentiation

To induce neural differentiation, iPSCs were plated under feeder-free conditions described above. Twenty-four hours after plating, media was changed to either the “Dorsal” condition (mTESR1 plus 100nM LDN193198 and 2uM SB431542) or the “Accelerated Dorsal” condition (Stem Cell Technologies AggreWell medium plus 100 nM LDN193189 and 2 μM SB431542). Forty-eight hours later (Day 2), the Dorsal condition media was replaced and the “Accelerated Dorsal” condition media was changed to N2/B27 medium plus 100nM LDN193189 plus 2uM SB431542. On Days 4 and 6, media for both conditions was replaced. Total RNA was collected (RNeasy Qiagen) under both conditions at Days 2 and 6, and at Day 4 for a subset of samples. The Accelerated Dorsal Condition was exclusively used for continued neural differentiation and media was replaced daily up until Day 9. On Day 9, Total RNA was collected or cells were passaged using Accutase onto Poly-L-ornithine\fibronectin coated tissue culture dishes in N2 media plus 20ng\ml FGF2. Media was changed each day up until Day 15 when RNA was harvested or cells were passaged with HBSS onto Poly-L-ornithine\fibronectin coated tissue culture dishes in N2 media plus 20ng/ml FGF2. Media was exchanged each day with fresh N2 until Day 21 when neural rosettes appear. On Day 21, Total RNA was collected or cells were passaged with HBSS onto PDL\Laminin coated coverglass with or without Rat E18 astrocytes in N2 media in a humidified 37°C tissue culture incubator at 5% oxygen. After 24 hours, the media was exchanged for Neuronal Differentiation Media (NeuroBasal Invitrogen 12348-017, 1X GlutaMax Invitrogen 35050-061,. 3nM Selenite, 25ug/ml Insulin Sigma I6634, 1X Pen/Strep final 1X Invitrogen 15140-122, 1X B27 Invitrogen 17504-044, concentration, 10ng/ml BDNF R & D systems. 248-BD/CF and 10ng/ml NT3 R & D systems, 267-N3/CF). 50% of the media was changed every other day until Day 28. After seven days on astrocytes, 100% of the media was changed to Neuronal Differentiation Media plus 20uM AraC. 50% of the media was changed with Neuronal Differentiation Media plus 20uM AraC every other day up to Day 35. At Day 35, 100% of the media was exchanged for Neuronal Differentiation Media without AraC and 50% of media was exchanged every other day for up to 8 weeks. RNA was harvested at indicated intervals during the process.

### Electrophysiology

Human neuronal cultures on glass coverslips were submerged in our recording chamber and constantly perfused with an external bath solution consisting of (in mM): 128 NaCl, 30 glucose, 25 HEPES, 5 KCl, 2 CaCl2 and 1 MgCl2 adjusted to pH 7.35 with NaOH. All recordings were performed at approximately 32°C. Patch pipettes were fabricated from borosilicate glass (N51A, King Precision Glass, Inc.) to a resistance of 2-5 MΩ. For voltage-clamp measurements, cells were held at -70mV and recording pipettes were filled with (in mM): 125 potassium gluconate, 10 HEPES, 4 Mg-ATP, 0.3 Na-GTP, 0.1 EGTA, 10 phosophocreatine, 0.05%, adjusted to pH 7.3 with KOH. Current signals were recorded with either an Axopatch 200B (Molecular Devices) or a Multiclamp700A amplifier (Molecular Devices) and were filtered at 2 kHz using a built in Bessel filter and digitized at 10 kHz. Data were acquired using Axograph on a Dell PC (Windows 7). We tested for differences in capacitance and membrane resistance across time in culture using linear mixed effects modeling, treating line and donor as random intercepts, and natural log-transforming these two measures to improve normality assumptions of these models.

### Cellular Imaging

Neural progenitor cells spontaneously self-organize into rosette structures reflecting morphological properties of the developing neural tube (*42*). Neural progenitors were plated in 24 well ibidi plates in triplicate for each line. After differentiation for 6 days, an acellular lumen is detectable with antibody directed against ZO-1. Surrounding the lumen are laminae of dorsal forebrain progenitors identified with anti-OTX2 antibodies. Nuclei were labeled with DAPI. Under these conditions, the structures are largely 2-dimensional but are becoming pseudo-3D. An array of 6×6 widefield (non-confocal) images was captured automatically from each of three neighboring wells using the Operetta and processed using custom code in Columbus (both Perkin Elmer). Briefly, ZO-1 segmentation was used to reveal the core of each rosette. Anti-OTX2 immunoreactivity was used to identify the limits of each rosette. Nuclear morphology was measured indirectly using the DAPI signal.

## RNA sequencing (RNA-seq) data generation, processing and quality control

### RNA sequencing

Total RNA was extracted from samples using the RNeasy Plus Mini Kit (Qiagen). Paired-end strand-specific sequencing libraries were prepared from 300ng total RNA using the TruSeq Stranded Total RNA Library Preparation kit with Ribo-Zero Gold ribosomal RNA depletion (Illumina). An equivalent amount of synthetic External RNA Controls Consortium (ERCC) RNA Mix 1 (Thermo Fisher Scientific) was spiked into each sample for quality control purposes. The libraries were sequenced on an Illumina HiSeq 3000 at the LIBD Sequencing Facility, after which the Illumina Real Time Analysis (RTA) module was used to perform image analysis and base calling and the BCL converter (CASAVA v1.8.2) was used to generate sequence reads, producing a mean of 58.3 million 100-bp paired-end reads per sample.

### Processing Pipeline

Raw sequencing reads were mapped to the hg38/GRCh38 human reference genome with splice-aware aligner HISAT2 version 2.0.4 (*43*). Samples without rodent tissue averaged 86.6% alignment rate (SD=5.1%) while 31 samples co-cultured with rat astrocytes averaged 33.9% alignment rate (SD=10.1%) to the human genome. Feature-level quantification based on GENCODE release 25 (GRCh38.p7) annotation was run on aligned reads using featureCounts (subread version 1.5.0-p3) (*44*) with a mean 62.9% (SD=8.2%) of mapped reads assigned to genes for human samples, and a mean 43.1% (SD=9.4%) of mapped reads assigned to genes for samples containing rat astrocytes. Exon-exon junction counts were extracted from the BAM files using regtools (*45*) v. 0.1.0 and the ‘bed_to_juncs’ program from TopHat2 (*46*) to retain the number of supporting reads (in addition to returning the coordinates of the spliced sequence, rather than the maximum fragment range) as described in (*26*). Annotated transcripts were quantified with Salmon version 0.7.2 (*47*) and the synthetic ERCC transcripts were quantified with Kallisto version 0.43.0 (*48*). For an additional QC check of sample labeling, variant calling on 740 common missense SNVs was performed on each sample using bcftools version 1.2.

### Quality control

After pre-processing, samples were checked for quality control measures. All samples passed QC checks for alignment rate, gene assignment rate, mitochondrial mapping rates, and ERCC spike-in concentrations. We looked for batch effects by checking for differences in technical metrics by batch, or for separation by batch within top principal components. Additionally, we clustered the gene expression of three sets of replicates sequenced on seven different flow cells, finding that the replicates clustered by sample and not flow cell. All of our batch effect checks indicated consistency across batch. Next we examined expression through the differentiation time-course of known pluripotency and neuronal differentiation marker genes to confirm our cell lines differentiated as expected; one cell line (six samples) did not pass this marker check due to slow differentiation and was dropped from all analyses. Finally, a genotype check was conducted to confirm the donor labeling of all samples. Called coding variants were matched to existing microarray-based genotype calls, and five samples were dropped due to ambiguous sample identity.

### Rat astrocytes

Purified rat astrocytes from five samples, as well as the co-cultured neuron/astrocyte samples and four samples of human neurons, were processed with an analogous rat pipeline as above (see *Processing Pipeline*). The samples were aligned to the rn6/Rnor_6.0 genome and feature counts were quantified using the Ensembl release 86 annotation of the rat transcriptome. A mean 90.7% (SD=2.1%) of rat astrocyte reads aligned to the reference genome and co-cultured samples had an average 68.4% (SD=9.9%) alignment rate, while human neurons averaged 16.7% (SD=4.5%) alignment. The expression levels of the three groups were compared to each other in assessing the neuronal maturation effects of co-culturing with astrocytes.

### Public data processing

Multiple public human RNA-seq datasets were downloaded and used to quantify the developmental stage and cellular composition of our samples across the differentiation time-course. All public data below were run through the same processing pipeline as outlined above to get comparable read counts and expression values (see *Processing Pipeline*).

Raw FASTQ files from each of the following datasets were downloaded from the sequencing read archive (SRA).

1. CORTECON (van de Leemput et al. (*7*)) included 24 single-end (50bp) samples of human cerebral cortex development from hESCs across nine timepoints (days 0, 7, 12, 19, 26, 33, 49, 63, and 77) [SRP041179, GSE56796];
2. The ScoreCard dataset (Choi et al. (*49*), Tsankov et al. (*15*)) consisted of 73 paired-end (2×100bp) samples of hESCs, hiPSCs, and fibroblasts [SRP063867, GSE73211];
3. Song et al. (*22*) consisted of 174 single cell samples of iPSCs, NPCs, and motor neurons from both paired- and single-end libraries [SRP082522, GSE85908];
4. Close et al. (*8*) included 1733 single cell and 40 pooled samples of paired-end (2×50bp), hESC-derived cortical interneurons at four timepoints of differentiation (days 26, 54, 100, and 125) profiling both neurons (DCX+) and progenitors (DCX-) [SRP096727, GSE93593];
5. Darmanis et al. (*23*) included 420 paired-end (2×75bp) single cell samples of four embryonic and eight adult brains from eight types of cortical tissue [SRP057196, GSE67835].
6. Bardy et al. (*34*) consisted of 56 paired-end (2×100bp) single cell samples of iPSC-derived neurons, as well as corresponding Patch-seq electrophysiology data.
7. Tekin et al. (*32*) included single cell paired-end reads of induced neurons both on and off human and mouse astrocytes at 1 week and 5 weeks after differentiation. Additionally, FASTQ files from both bulk and single cell RNA-seq datasets were downloaded from the BrainSpan atlas (*30*).
8. Homogenate data consisted of 407 single-end (75bp) samples from neocortical regions of 41 donors aged early fetal through adult (40 year old);
9. Single cell data were 932 single-end (100bp) samples from the DLPFC and dorsal pallium of eight early- to mid-fetal brains. The following data from the BrainSeq Phase I consortium was downloaded and reprocessed (*26*).
10. The BrainSeq data consisted of one DLPFC sample each from 318 unique donors ranging from fetal through age 85, with paired-end (2×100bp) libraries. Other datasets used for evaluating the cellular deconvolution approaches included:
11. Raw FASTQ files from human organoids (both bulk and single cell data) from Sloan et al. (*33*), which were processed with the above pipeline [SRR5676732];
12. Publicly available gene counts (not FASTQ files) for Ensembl v70 from Hoffman et al (*31*).

## Statistical Analyses *WGCNA*

To identify dynamic patterns of gene expression across neuron maturation, we performed signed weighted gene co-expression network analysis (WGCNA) (*16*) using the software’s R package. The analysis was performed on 25,466 expressed genes (cutoff mean RPKM > 0.1) from 106 samples across all time points (days 2, 4, 6, 9, 15, 21, 49, 63, and 77). Self-renewal samples and neurons cultured without rat astrocytes were not included in the WGCNA analysis. Normalized expression values in the form of log_2_(RPKM+1) were used, with gene assignment rate (as a measure of sample quality) regressed out of the expression matrix. To find clusters, the software first selected a soft thresholding power of six, then, using a minimum module size of 30 and maximum block size of 10000, assigned 22,182 of the expressed genes to eleven signed modules representing dynamic expression patterns through differentiation. GO analysis was then carried out on the modules to find biological processes and functions enriched by the gene sets of each cluster. To evaluate replication of the expression patterns, we separated the reprocessed CORTECON data into the same eleven gene sets, and compared both the eigengenes and GO terms of the eleven modules calculated in each of the two datasets.

### Time-course DE

Again looking at 106 samples and 25,466 expressed genes, we performed differential expression analysis to find genes with changing expression through the stages of neuronal differentiation. Our statistical model used the voom method (*50*) of linear modeling to estimate the mean-variance relationship of the gene log-counts, adjusting for cell line and the proportion of mapped reads assigned to genes (gene assignment rate). We found genes most differentially expressed (DE) between each of the neighboring cell conditions of early neuronal differentiation, NPC, rosettes, and neurons. Within the same model we also found genes differentially expressed across the entire differentiation time-course. Voom modeling was run on genes, exons, exon-exon junctions, and annotated transcripts, with p-values adjusted for false discovery rate (FDR) within feature type.

### Alternative splicing

Alternative splicing events were further investigated through intron retention (IR) measures and isoform shifts. Ratios of retained introns to spliced introns were calculated using IRFinder version 1.1.1 (*51*), and linear regression models were used to obtain a list of genes with significantly increasing or decreasing IR ratios across the time-course, adjusting for cell line and gene assignment rate. GO analysis on the directional gene lists was then completed. Additionally, the percent of aligned reads assigned to introns was calculated by featureCounts for each sample using a GTF of the intronic features of GENCODE release 25. Differences in intron assignment rate by condition was evaluated with a linear regression model adjusting for cell line. Isoform shifts were determined using the transcript quantification output by the processing pipeline described previously. Normalized transcript expression levels (TPMs) were averaged across samples grouped by each of the four major cell conditions (accelerated dorsal, NPC, rosette, and differentiated neurons). Genes with more than one highest-expressing transcript across the four conditions were considered to have an isoform switch. Major isoforms with lengths shorter than 250bp were not considered to be switching events.

### Astrocyte Effects

To assess the effect of astrocytes on neuronal maturation, we compared the gene expression at day 77 of four neuronal lines cultured alone to seven of the same lines co-cultured with rodent astrocytes. A voom model adjusting for cell line and gene assignment rate was implemented to find genes significantly differentially expressed between the two groups at FDR<0.05. We then performed GO analysis on the up-regulated and down-regulated gene sets to investigate enrichment of the DE genes. Similarly, using a voom model adjusting for day and gene assignment rate, we tested for differential expression of rat genes between the co-cultured samples and three purified rat astrocytes on day 49, 63, and 77, followed by GO analysis on the up- and down-regulated genes.

### Public data

We used two regression calibration models to determine the relative compositions of our iPSC model system. The first model involved identifying the relative developmental stage of our sequenced cells, using data from the ScoreCard (*15, 49*) and BrainSpan (*30*) homogenate sequencing projects. The second model involved identifying the relative cellular composition of our sequenced cells, using the Fluidigm-based single cell RNA-seq datasets from Song et al. (*22*) and Darmanis et al. (*23*).

Developmental stage model: we combined normalized expression data (log2(RPKM+1)) from the 21 iPSC samples in the ScoreCard project and 407 neocortical bulk samples from the BrainSpan project across seven timepoints (73 early-, 73 mid- and 17 late-prenatal, as well as 53 infant, 71 child, 55 teen and 65 adult postnatal). We defined stage-specific genes using the same framework described by Jaffe and Irizarry 2014 (*24*), which involved creating a “barcode” of 25 genes per stage, that were more highly expressed for each stage compared to all others (t-statistic p-values < 1e-15), and ranking subsequent significant genes by log2 fold changes for selection. As some stages had similar expression levels with others, we ended up with 169 unique genes and created the regression calibration design matrix based on Houseman et al (*25*), shown in Table S8. We then projected samples into this design matrix using the ‘projectCellType()’ function in the *minfi* Bioconductor package (*52*).

Cell proportion model: we combined single cell normalized expression data (log2(RPKM+1)) from 63 iPSC and 73 NPC samples from Song et al. (*22*), and 25 replicating and 110 quiescent fetal neurons, 18 oligodendrocyte progenitor cells (OPCs), 131 neurons, 62 astrocytes, 38 oligodendrocytes,16 microglia and 20 endothelial cells from Darmanis et al (*23*). We defined cell type-specific genes using the same framework as the developmental stage model. Here we created a “barcode” of 25 genes per cell type that were more highly expressed for one cell type compared to all others (t-statistic p-values < 1e-15), and we ranked by log_2_ fold changes for selection. From our final set of 228 unique genes, we scaled each gene expression value to the standard normal distribution to improve comparability between single cell and bulk RNA-seq data, and created the regression calibration design matrix based on Houseman et al. (*25*), shown in Table S9. We again then projected samples into the design matrix using the ‘projectCellType()’ function in the *minfi* Bioconductor package. Forty-four genes were shared between these two statistical models for deconvolution.

## Data and Software availability

Accompanying processing and analysis code is available at: https://github.com/lieberinstitute/libd_stem_timecourse and all raw sequencing reads are available through SRA at the following accession: [TBD].

## Supplementary Figures

**Figure S1:**
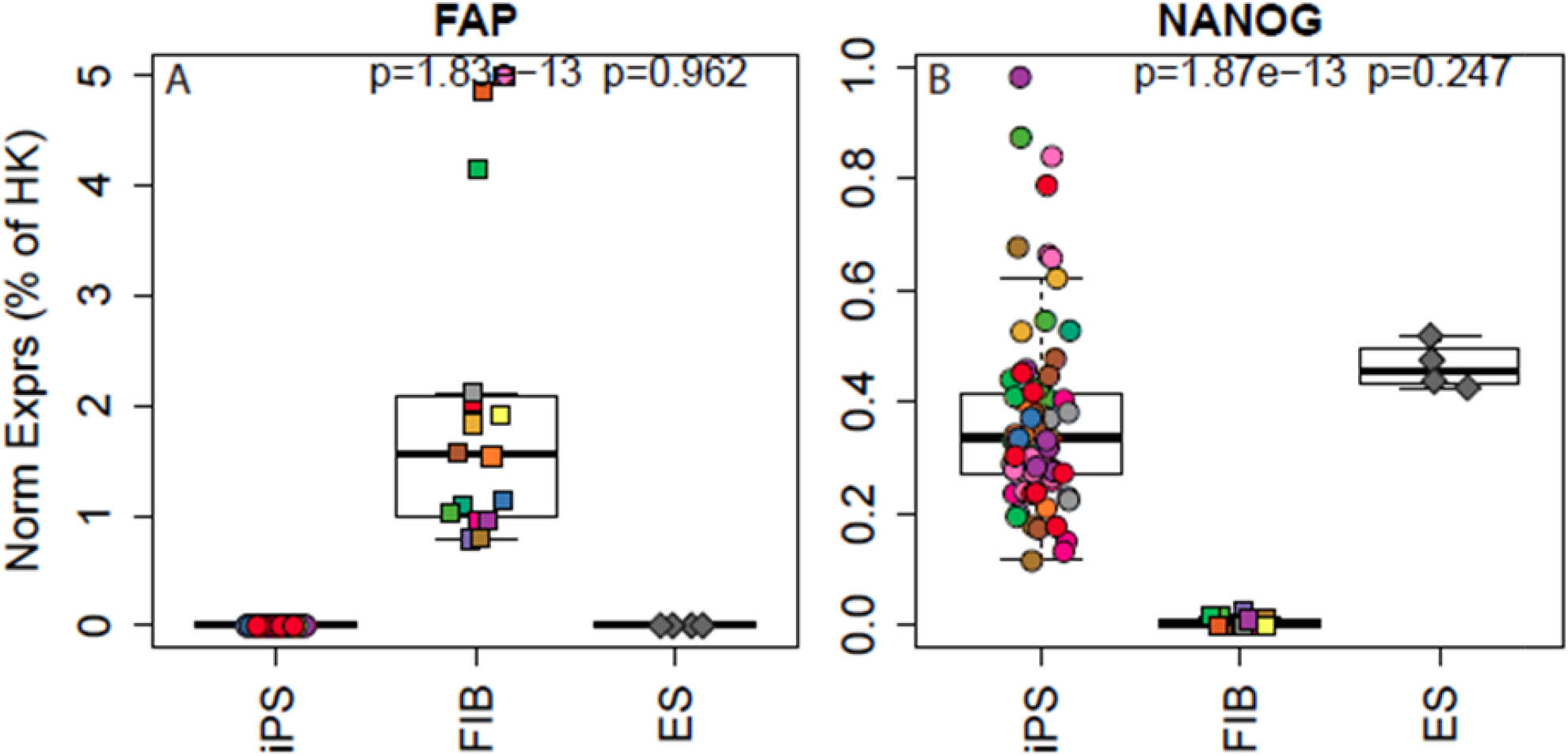
Fluidigm-based expression profiling. Fluidigm-based expression profiling confirmed the loss of *FAP* (Fibroblast Activation Protein Alpha) expression across all iPSC lines and the gain of *NANOG* expression.

**Figure S2:**
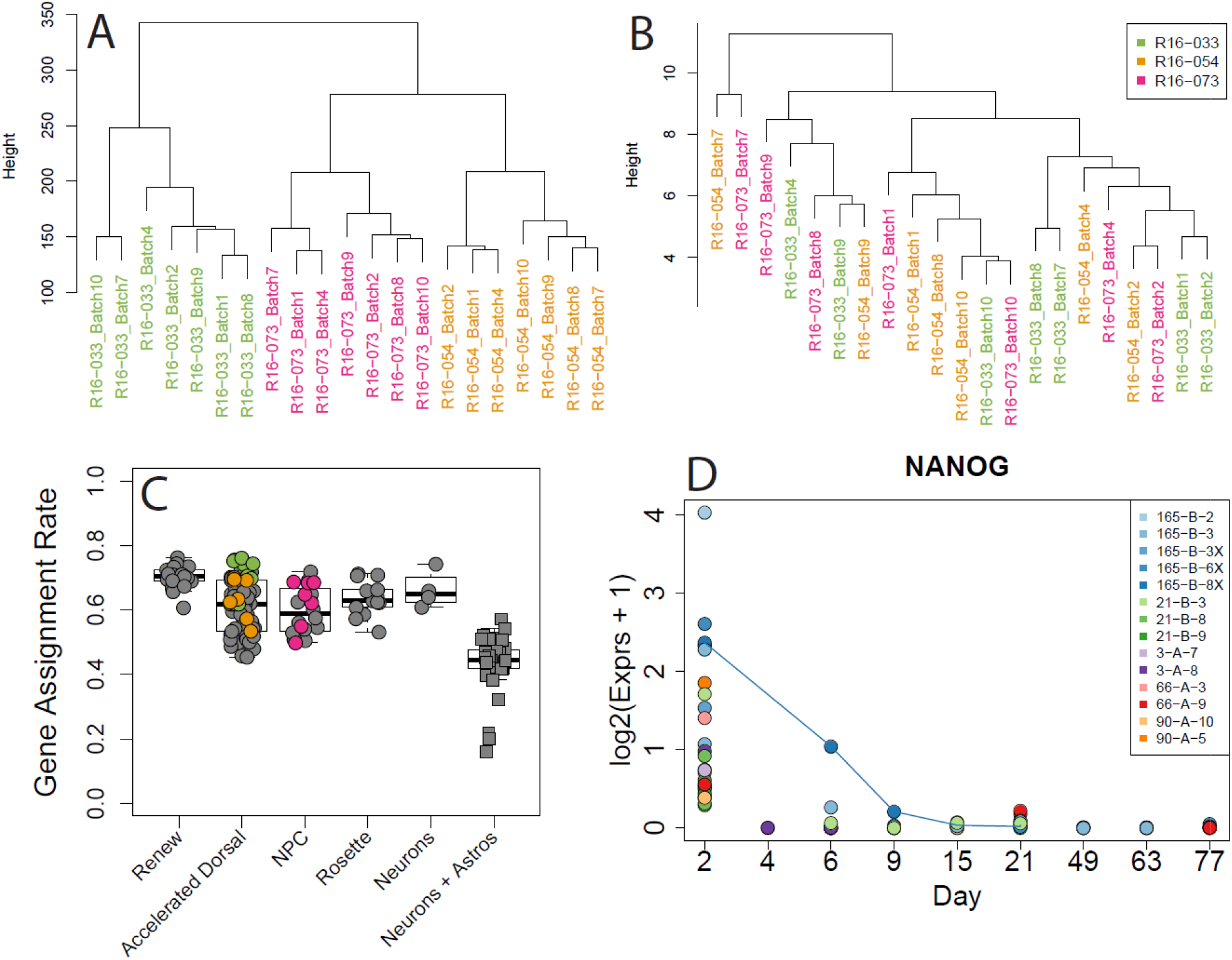
RNA-seq quality control. (A) Gene expression levels of three RNA replicates across seven flow cells clustered by cell line, and (B) ERCC spike-ins of the replicates clustered more by flow cell (batch number). (C) The percent of aligned reads assigned to genes across cellular conditions, with colored replicate points having similar rates. (D) Slow differentiation of cell line 165-B-8X as shown by expression of *NANOG* through time, resulting in us dropping the 6 samples from this line.

**Figure S3:**
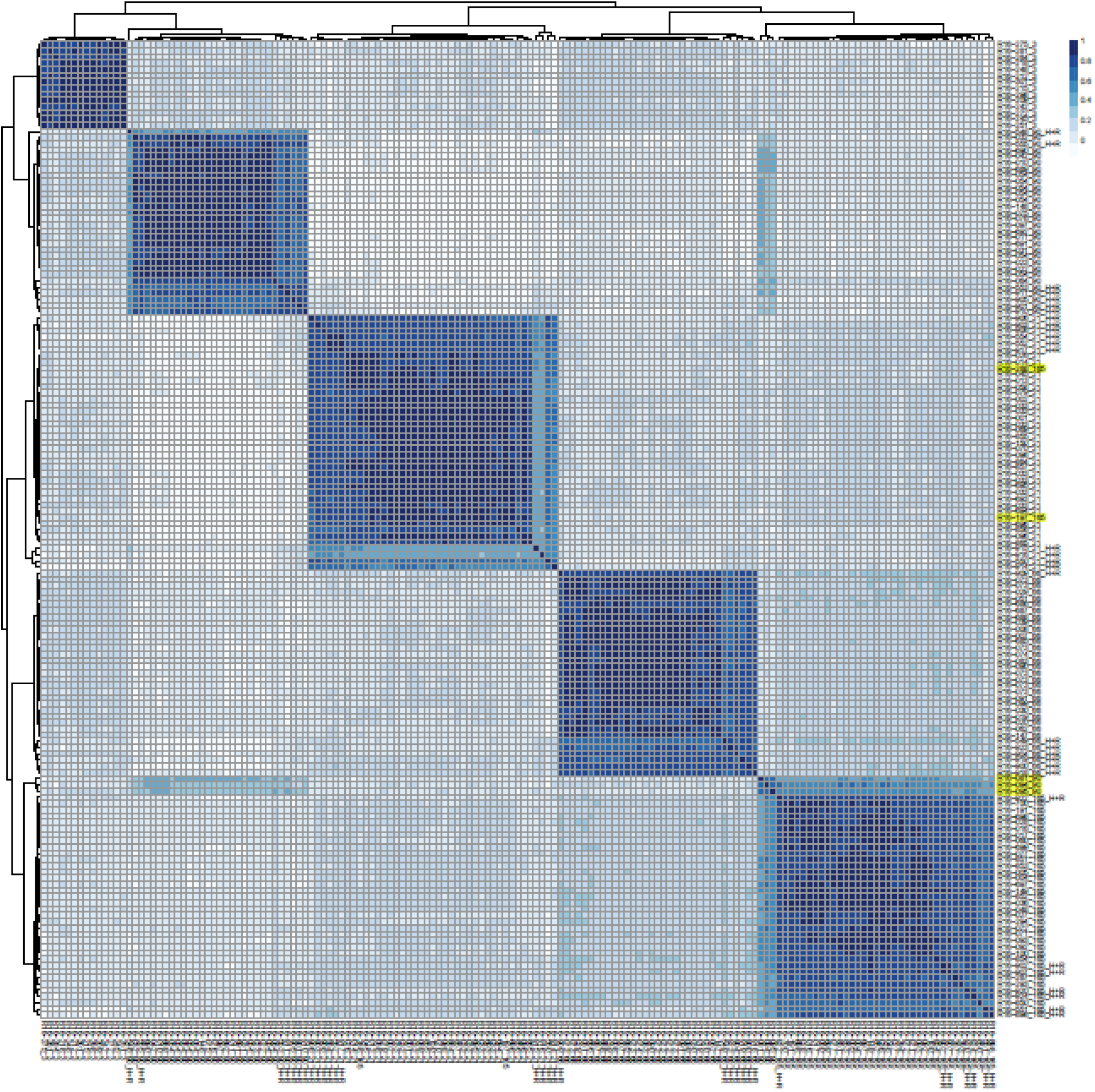
Genotyping mismatches. Correlation heatmap of microarray-based genotype calls using called coding variants. Five samples (highlighted in yellow) did not highly correlate with their labeled donor identity and were dropped from all analyses.

**Figure S4:**
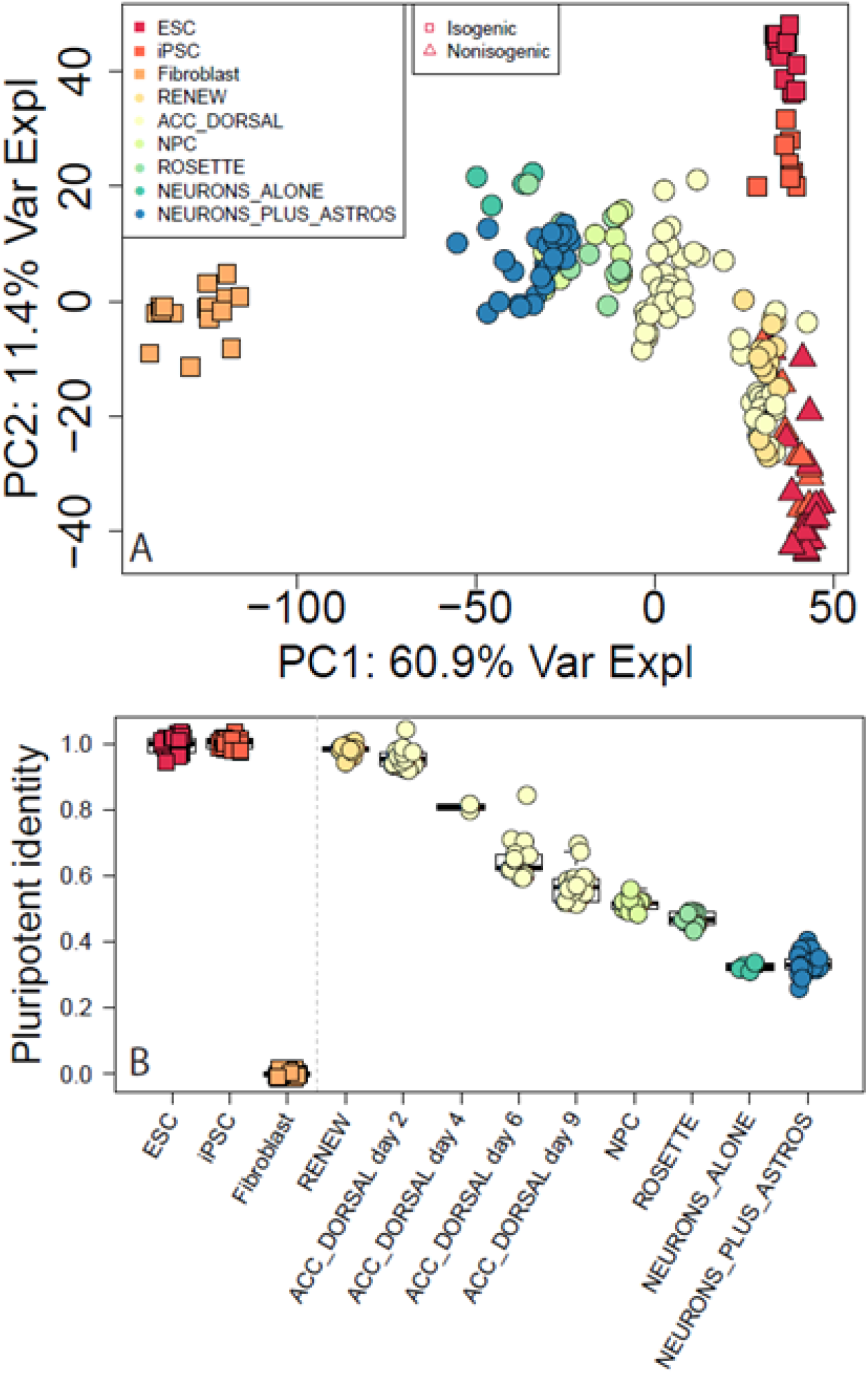
Comparison to ScoreCard data. (A) Our data (circles) projected into the first two PCs calculated from the ScoreCard reference data gene expression levels (square and triangle points), to assess the representativeness of our iPSC cell lines and subsequent differentiation data. (B) The pluripotent identity of ScoreCard data and our data, showing our self-renewal/iPSC lines with a mean 98.1% pluripotency identity which significantly decreased through differentiation.

**Figure S5:**
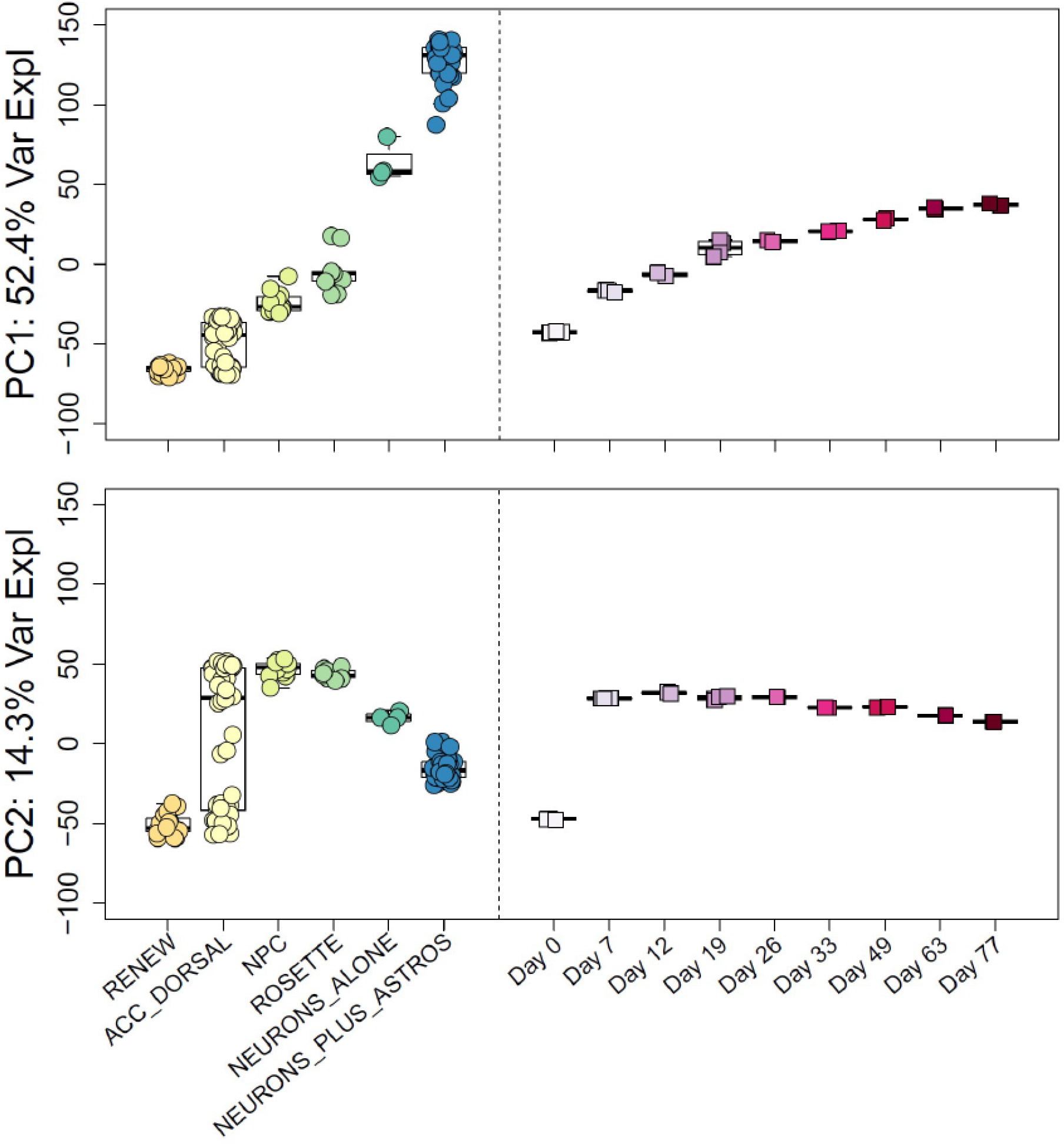
PCA and comparison to CORTECON data. PCA of gene expression levels showing PC1 representing corticogenesis, and PC2 separating NPC stage cells from early days as well as mature neurons. Both of these components of variability — a linear trend in PC1 across differentiation and a quadratic trend in PC2 — were significantly conserved when we projected the CORTECON data (square points) into these PCs.

**Figure S6:**
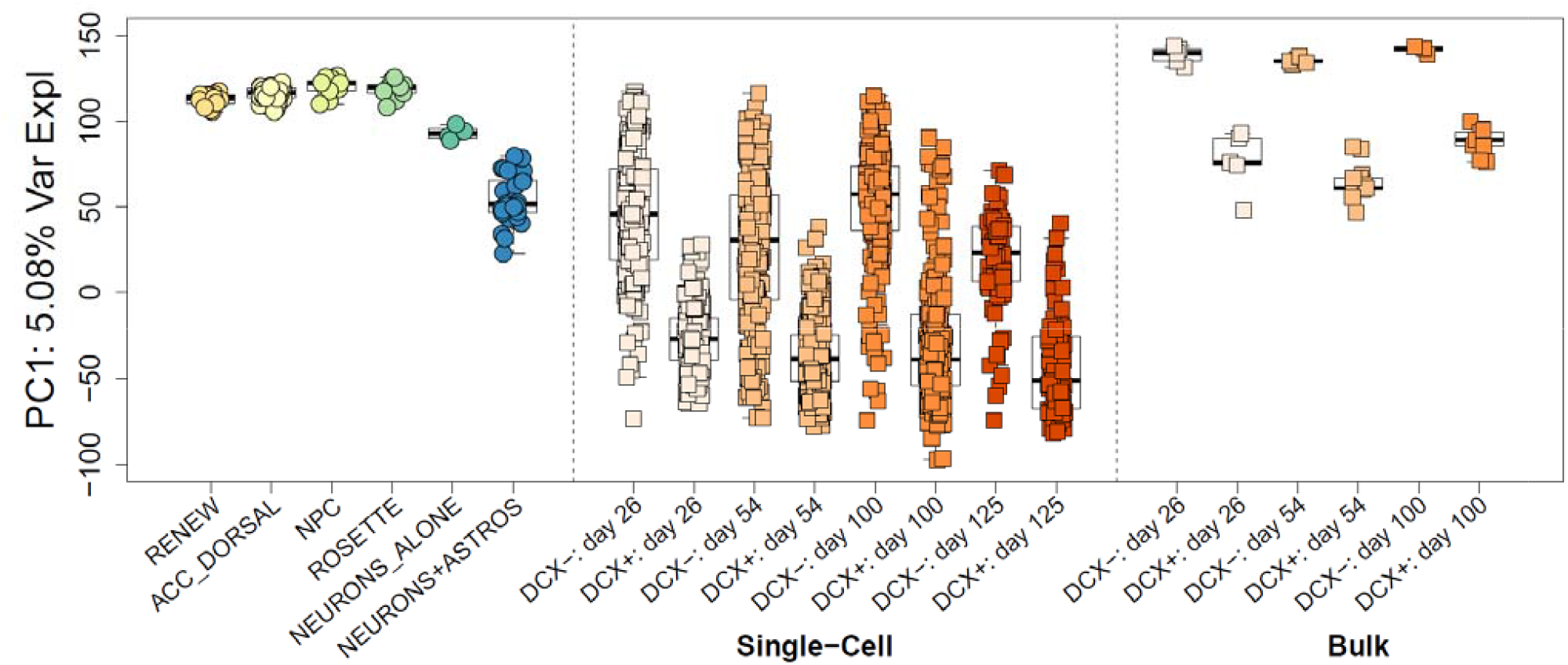
PCA of Close et al. PCs based on pooled cell-level data from Close et al., with our data projected in. Our cells clustered with the Close et al. pooled cells, with our more mature neuronal lines clustering with DCX+ samples and early NPC lines clustering with DCX-samples.

**Figure S7:**
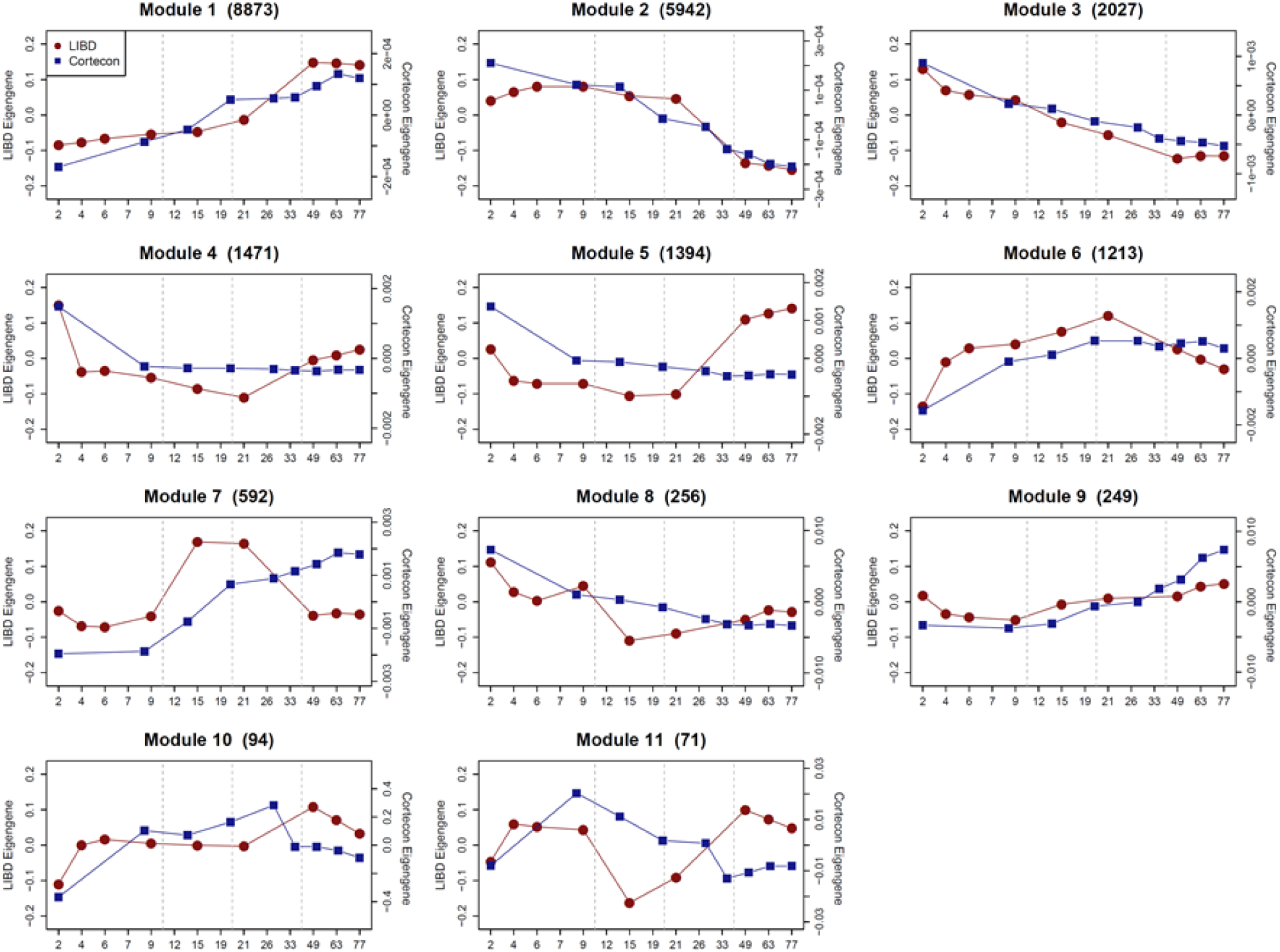
Eigengenes of WGCNA modules. The eigengenes of the WGCNA modules calculated from our data, plotted with the CORTECON eigengenes calculated after separating that data into the same eleven gene sets, showing similar developmental patterns across most of the top modules.

**Figure S8:**
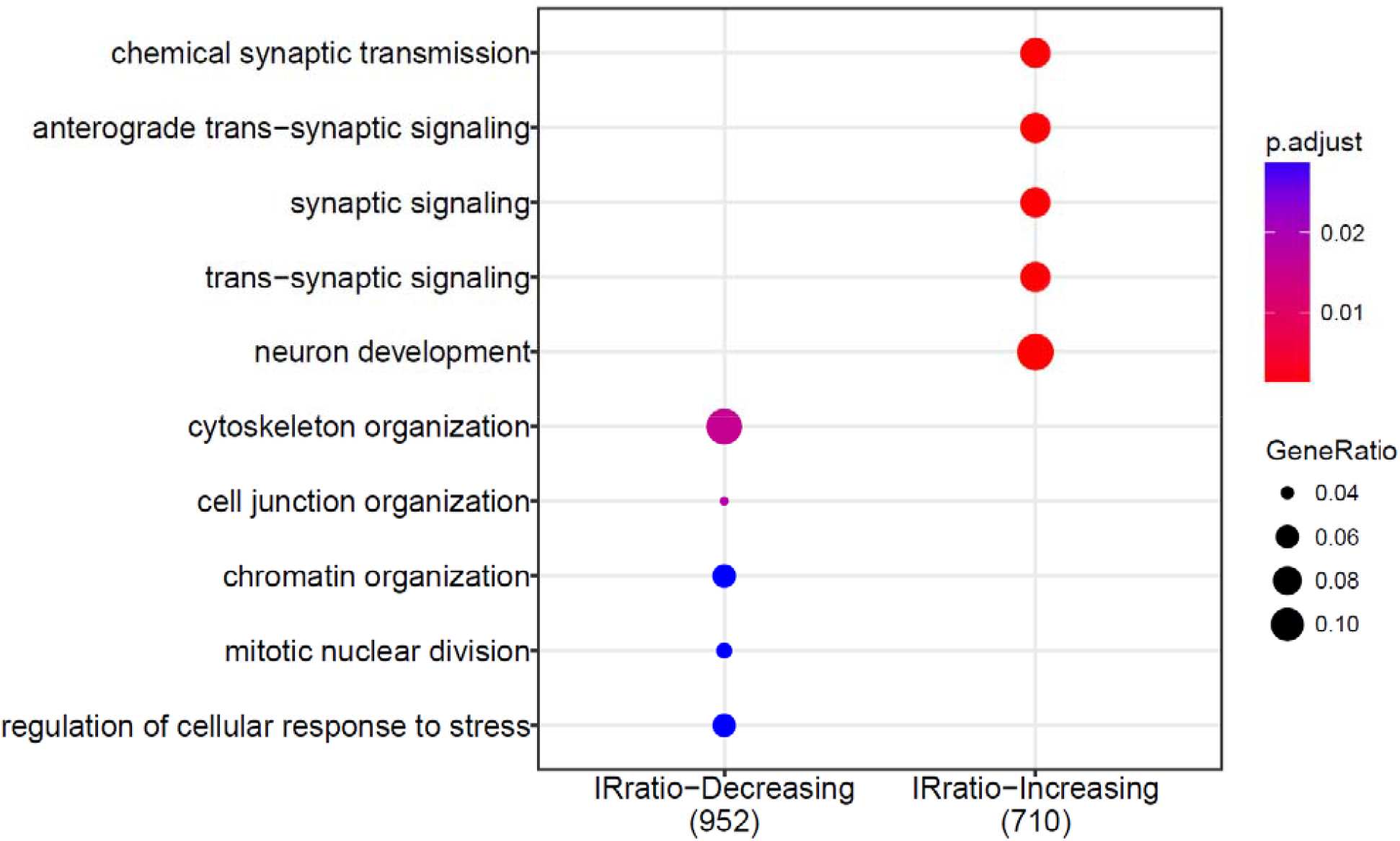
GO analysis of intron retention ratios. Enrichment of the 2847 genes with significantly (FDR < 0.1%) decreasing (1518/2847) or increasing (1329/2847) intron retention ratios. As expected, increasing intron retention was enriched for neuron development terms.

**Figure S9:**
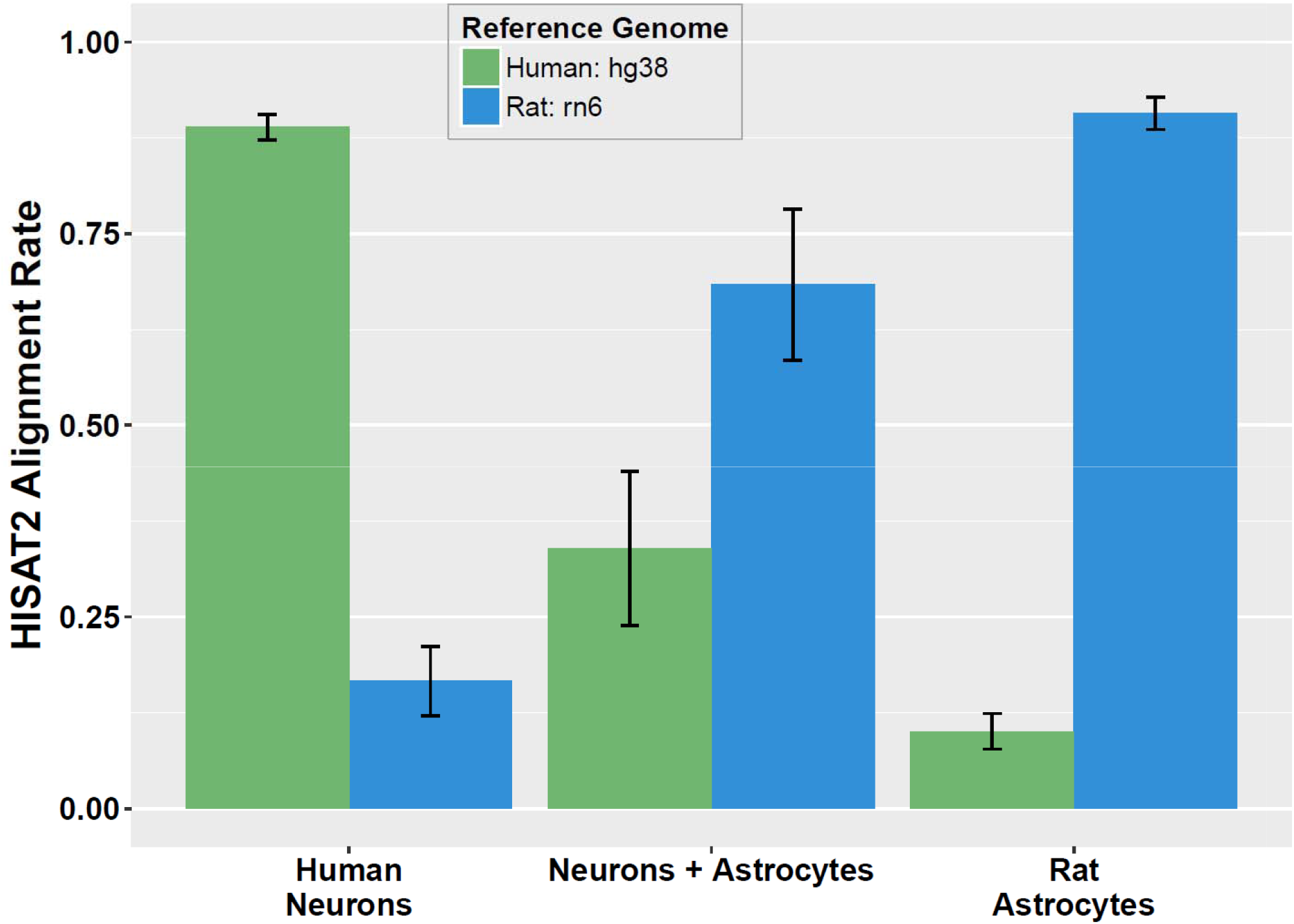
Cross-species mapping in RNA-seq alignments. Alignment rates of human neurons alone, rat astrocytes alone, and co-cultured samples mapped to human and rat genomes. Human neuron samples had a low alignment rate to the rat genome (mean=16.6%) and rat astrocytes lowly aligned to the human genome (mean=10.1%). We took these two sets of aligned reads and mapped them back to human and rat, respectively, and found that in each case only 3 genes accounted for the majority of cross-mapped expression (Table S5).

**Figure S10: Cell type deconvolution proportions of 131 signature genes**. Boxplots of the standardized expression levels of all 131 genes that distinguish iPSCs, NPCs, fetal replicating neurons, fetal quiescent neurons, adult neurons, and adult endothelial cells.

See: Figure_S10_markerExprsZ_boxplots.pdf

**Figure S11:**
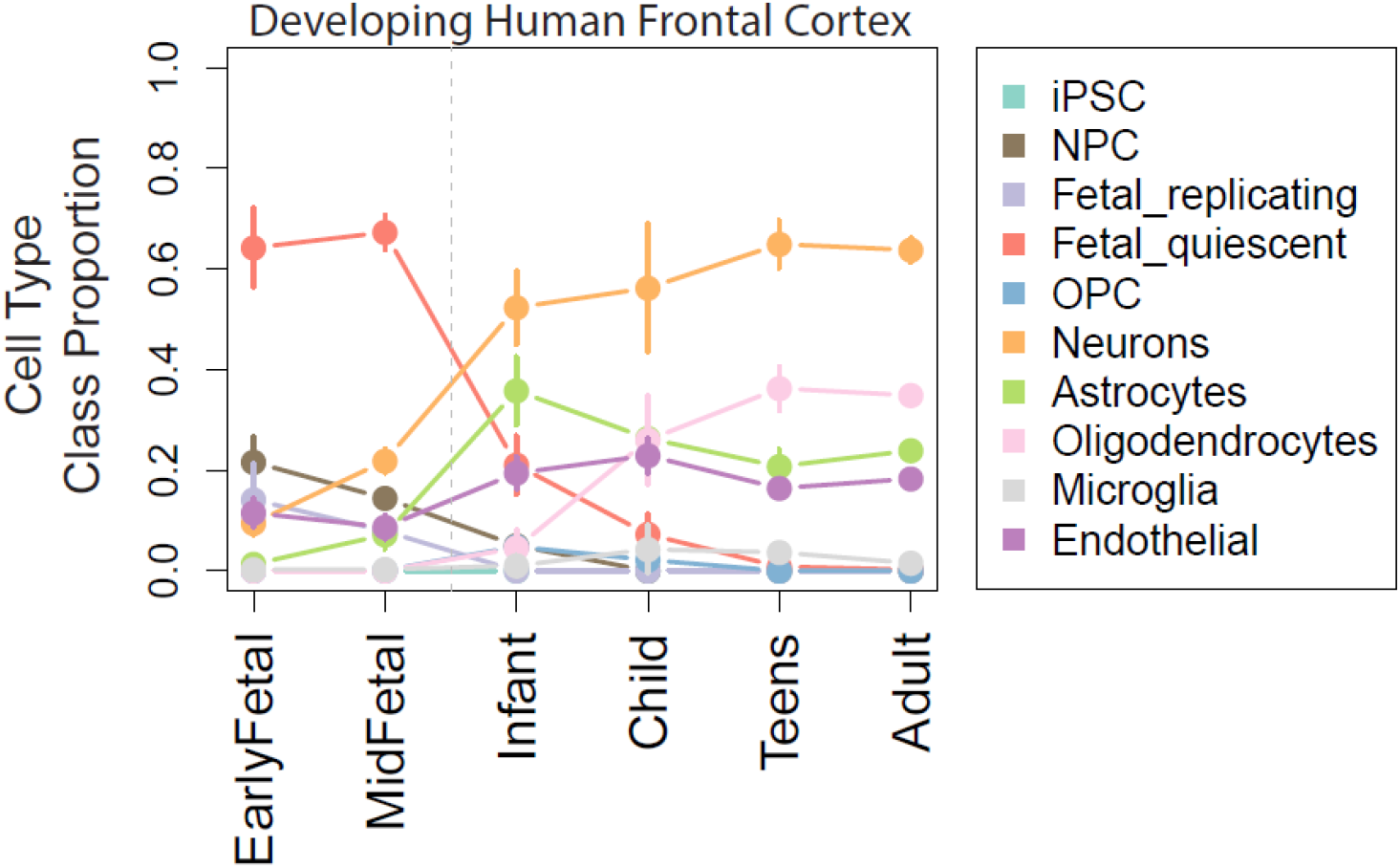
Deconvolution on postmortem brain tissue. We applied our cell type RNA deconvolution approach to a large RNA-seq dataset from postmortem human brain tissue and observed the expected loss of fetal quiescent neurons and NPCs, and rise of adult neurons and oligodendrocytes.

**Figure S12:**
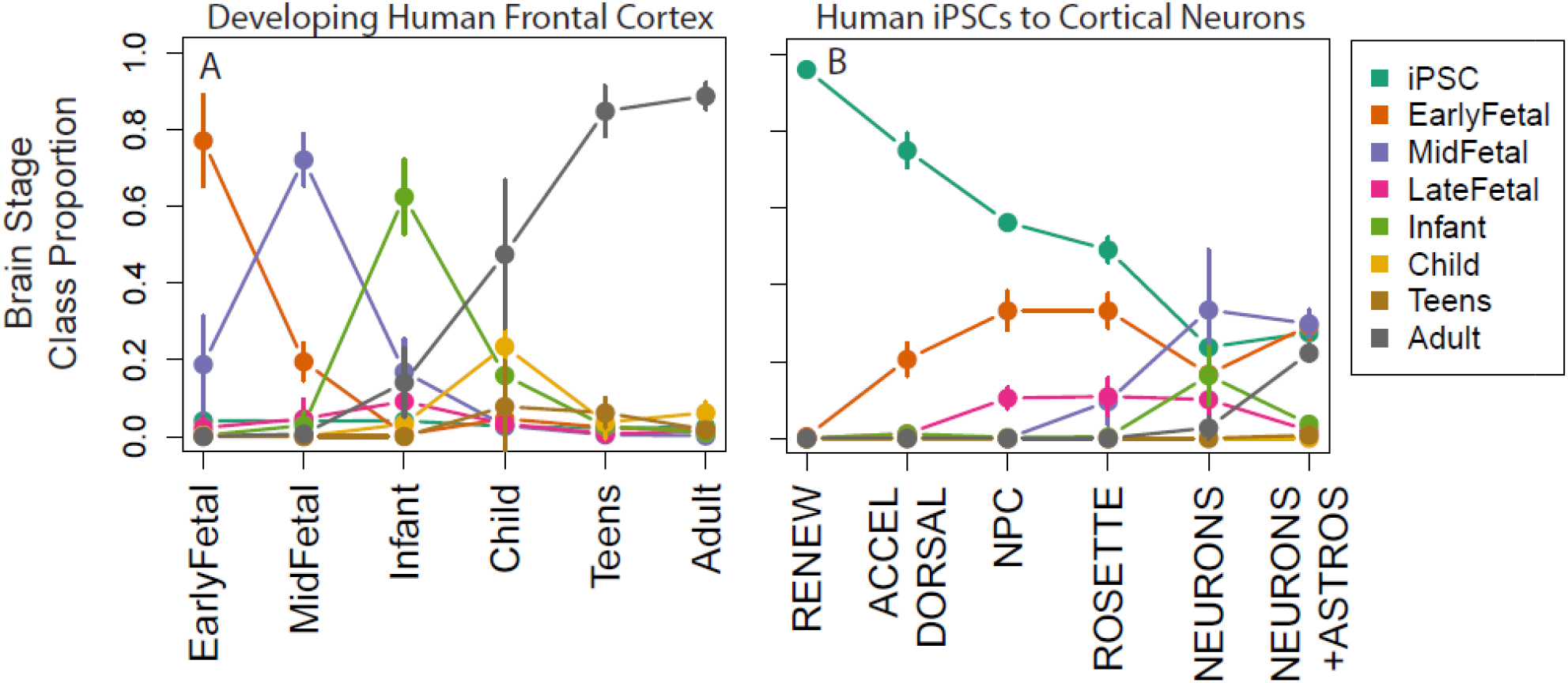
Brain stage deconvolution. We designed a second deconvolution model to estimate RNA fractions from eight developmental stages. We applied the deconvolution to a large postmortem brain tissue dataset (A) and our stem cell data (B). We observed a general loss of pluripotency through differentiation (p=3.77e-38) — the self renewing and early differentiating cells had high proportions of the iPSC signature (mean 96.1% and 75.1% respectively), with a rise of early-fetal neocortical-like signature (p=4.7e-7) first in the differentiating cells (20.6%) that became more prevalent in NPCs (33.3%) and rosettes (33.3%). We further identified the rise of a mid-fetal neocortical signature (p=3.0e-22) first appearing in rosettes (9.6%) and expanding into neurons off (33.5%) and on (29.9%) astrocytes. The most interesting class switch involved the late-fetal neocortical and adult neocortical classes — neurons grown off astrocytes had high class membership with late-fetal neocortex (10.1%) with low proportions of adult neocortex (2.7%). However, in the neurons co-cultured with rodent astrocytes, these proportions were reversed — these transcriptionally more mature neurons showed that 22.2% of RNA was analogous to adult neocortex and only 2.0% of RNA reflected late-fetal neocortex (Figure 5C).

**Figure S13:**
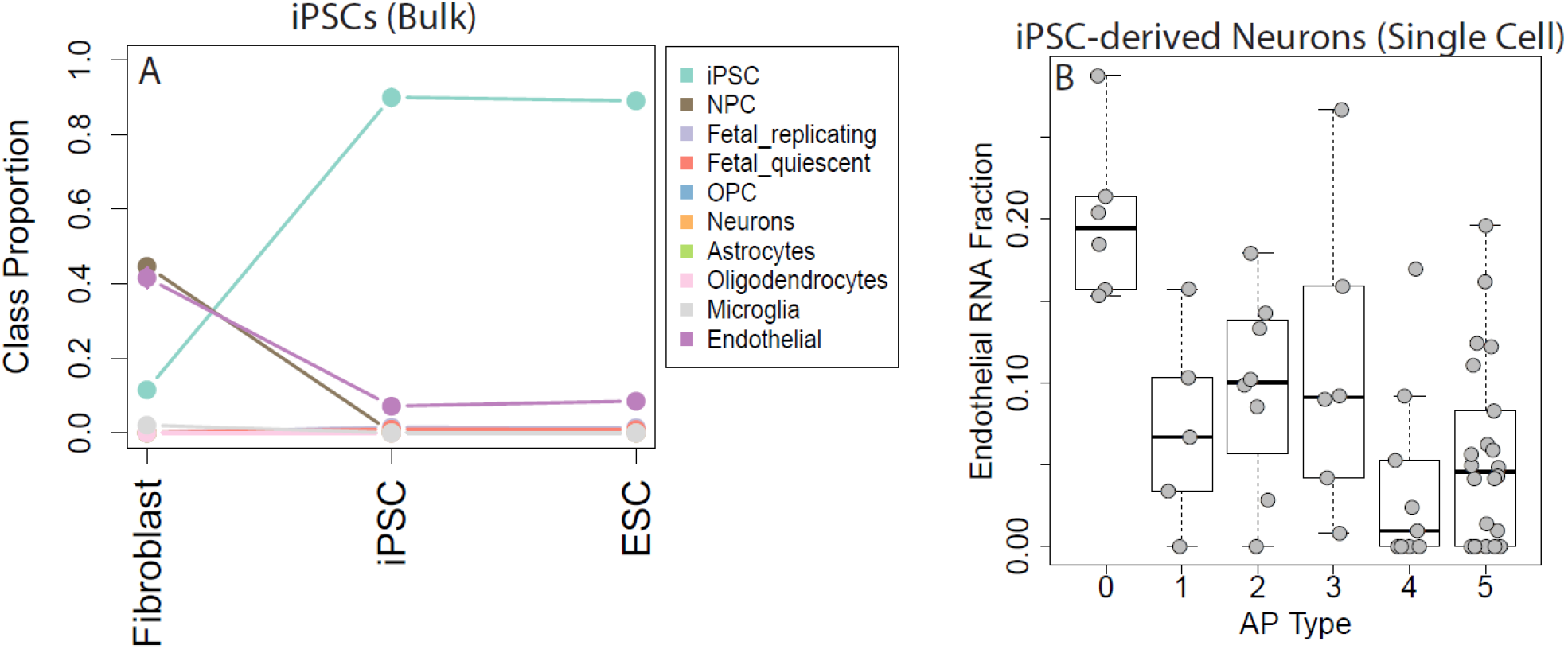
Deconvolution using public datasets. (A) The ScoreCard manuscript had reported a residual fibroblast-like signature which was not supported by applying this deconvolution to pure fibroblast and iPSC data. (B) A dataset of iPSC-derived neurons combined with activity state electrophysiology data suggest that the endothelial RNA fractions are associated with less mature astrocyte cells (Type 0, ~20% RNA Fraction).

### Supplementary Tables

**Table S1**: Gene set enrichment analysis of WGCNA modules.

**Table S2**: Gene and feature-level differential expression at FDR < 1%.

**Table S3**: Annotation classes of differentially expressed exon-exon junctions.

**Table S4**: Number of isoform shifts in differentially expressed genes.

**Table S5**: Details of the two sets of three highly expressed genes that contributed to the majority of reads that mapped across rat and human.

**Table S6**: List of 3214 genes differentially expressed (at FDR < 0.05) between the neurons alone and neurons on astrocytes.

**Table S7**: GO enrichment results of DE genes from on/off astrocyte analysis.

**Table 8**: 283 unique genes of the regression calibration design matrix for developmental stage model.

**Table S9**: 257 unique genes of the regression calibration design matrix for the cellular proportion model.

